# Non-Linear Dorsolateral Prefrontal Recruitment During Speech-in-Noise Perception in Aging

**DOI:** 10.64898/2026.08.14.744523

**Authors:** Maxime Perron, Frank A. Russo

**Affiliations:** Department of Psychology, Toronto Metropolitan University, Toronto, Ontario, Canada; Speech Language Pathology, University of Toronto, Toronto, Ontario, Canada

**Author notes:** Corresponding author: Maxime Perron, Department of Psychology, Toronto Metropolitan University, 350 Victoria Street, Toronto, Ontario, M5B 2K3, Canada.

**Keywords:** Neural Inefficiency, Speech-in-Noise, Listening Effort, Aging, Functional Near-Infrared Spectroscopy, Cognitive Compensation

## Abstract

Age-related speech-in-noise difficulties have been associated with increased activity in prefrontal regions. Whether this upregulation reflects adaptive compensation, neural inefficiency, or changes in task engagement as listening demands increase remains unresolved. Most studies have relied on linear contrasts of task difficulty, limiting our understanding of how the role of prefrontal recruitment evolves with listening demand. The present study examined these competing accounts within a non-linear framework. Using functional near-infrared spectroscopy, we measured activity in the dorsolateral prefrontal cortex (DLPFC) while 36 young and 34 older adults performed a sentence-in-noise task across five signal-to-noise ratio conditions. We tested whether DLPFC recruitment followed a quadratic trajectory across listening demands and whether its relationship with performance changed accordingly. Dorsomedial and ventrolateral prefrontal activity was also recorded to assess regional specificity. Bilateral DLPFC responses followed an inverted U-shaped pattern, increasing from easier to intermediate conditions before declining at the most difficult levels. Greater left DLPFC activation was associated with poorer performance in older adults across the ascending and peak portions of the demand-response function. High-performing older adults showed more youth-like recruitment profiles characterized by lower overall activation. Within the older adult group, higher left DLPFC activation was associated with greater age but not with hearing or global cognition. These results clarify the neural mechanisms underlying age-related speech-in-noise difficulties, suggesting that the shape of DLPFC recruitment reflects a demand-sensitive trajectory broadly preserved across age groups, while the level of activation in older adults may reflect neural processing efficiency, with greater upregulation associated with poorer performance.

## 1. Introduction

Speech-in-noise perception becomes increasingly difficult with age, and these difficulties cannot be explained fully by hearing loss or cognitive decline alone (Bilodeau-Mercure et al., 2015; Fostick et al., 2013; Fullgrabe et al., 2014). Age-related deafferentation may nevertheless contribute to these difficulties through downstream changes in central auditory processing. Reduced afferent input can trigger adaptive changes in central auditory processing, including increased central gain, which may partially restore neural response magnitude without restoring the fidelity of auditory representations (Bramhall and McMillan, 2024; Fabrizio-Stover et al., 2026). Together with age-related alterations in structural connectivity (Perron et al., 2021; Tremblay et al., 2019), such changes may increase the demands placed on higher-order systems during challenging speech perception. Consistent with this possibility, older adults frequently show greater recruitment of prefrontal cortex during effortful listening (Wong et al., 2009). However, the functional significance of this increased prefrontal recruitment remains uncertain.

Within the broader cognitive aging literature, increased prefrontal recruitment has been interpreted in two contrasting ways. Under a compensatory account, greater prefrontal activation reflects the adaptive mobilization of additional cognitive resources to counteract age-related sensory and neural decline and maintain performance (Cabeza et al., 2018; Park and Reuter-Lorenz, 2009). In contrast, a neural inefficiency account proposes that greater activation reflects less selective or less efficient recruitment and would therefore be expected to be associated with poorer rather than better performance (Logan et al., 2002; Reuter-Lorenz et al., 2001). Within the hearing literature, increased prefrontal recruitment in older adults has predominantly been interpreted through the compensatory lens (Du et al., 2016; Peelle, 2018; Wong et al., 2009). Compensation predicts that greater activation should accompany increasing listening demand and, critically, be associated with maintained or better behavioural performance.

A third possibility emerges when listening demands become extreme. Models of listening effort such as FUEL and MoLE propose that listeners may disengage when the perceived likelihood or value of successful comprehension no longer justifies continued effort (Herrmann and Johnsrude, 2020; Pichora-Fuller et al., 2016), predicting reduced neural recruitment and a decoupling of activation from performance at the most adverse levels of difficulty. Importantly, declining activation under these conditions would not necessarily indicate more efficient processing, but may instead reflect a reduction in the resources allocated to the task once continued effort is unlikely to yield sufficient benefit. Disengagement therefore provides a distinct account of reduced prefrontal activation at high levels of listening demand.

Distinguishing among these accounts therefore requires consideration not only of the magnitude of prefrontal recruitment, but also of its relationship with behavioural performance. Our previous fNIRS study revealed a pattern of DLPFC recruitment during speech-in-noise perception that was most closely aligned with the neural inefficiency account (Perron et al., 2025). Older adults showed greater bilateral DLPFC activation than younger adults under difficult listening conditions, yet greater activation was associated with poorer rather than better performance. Incorrect trials were also associated with greater DLPFC activation, and DLPFC upregulation negatively mediated the effect of age on accuracy. These findings suggested that increased DLPFC recruitment in older adults does not necessarily confer a behavioural benefit. However, that study sampled only a limited range of listening difficulty, precluding characterization of the full demand-response function and leaving unresolved whether the functional significance of DLPFC recruitment changes systematically as listening demands increase.

This question is important because effort-related responses are often nonlinear. Pupillometric, behavioural, and electrophysiological measures commonly increase as listening becomes more difficult before declining at the most adverse levels (Paul et al., 2021; Ryan et al., 2022; Wu et al., 2016; Zekveld and Kramer, 2014). Such inverted U-shaped functions have often been interpreted as reflecting progressive mobilization of cognitive resources followed by reduced resource allocation once listening demands become sufficiently adverse. Critically, however, the shape of this function alone does not establish the functional significance of the underlying neural recruitment. If the ascending portion reflects successful compensation, greater activation should predict better performance within that range. If increasing activation instead reflects inefficiency, greater activation should predict poorer performance. At the most difficult levels, disengagement should be reflected by declining activation together with a weakening or disappearance of the activation-performance relationship.

The present study therefore extends our previous work by providing a direct test of whether the functional significance of prefrontal recruitment changes across the listening-difficulty function. To do so, we sampled DLPFC responses across a substantially broader range of SNRs and tested whether the relationship between prefrontal recruitment and behavioural performance changed across that range. We predicted a nonlinear DLPFC response across SNR and tested whether brain-behaviour relationships across different levels of listening demand were consistent with compensation, inefficiency, or disengagement. We also examined whether these dynamics differed between high- and low-performing older adults, testing whether better speech-in-noise performance in aging is associated with preservation of a younger-like neural response profile.

Finally, we asked whether these effects were specific to the DLPFC or reflected a broader prefrontal response. We therefore recorded activity from the dorsomedial prefrontal cortex (DMPFC) and ventrolateral prefrontal cortex (VLPFC), regions with distinct functional roles in internally directed processing and speech-related predictive processing, respectively (Cope et al., 2017; Davis and Johnsrude, 2007; McKiernan et al., 2003). This regional comparison allowed us to determine whether demand-dependent changes in recruitment are specifically expressed within cognitive-control circuitry or distributed more broadly across prefrontal systems.

## 2. Method

### 2.1. Participants

Forty young adults and 39 older adults were recruited through emails, the Toronto Metropolitan University’s SONA participant pool, community advertisements, and word-of-mouth. Eligibility was verified using a Qualtrics online questionnaire (Qualtrics, Provo, UT). Eligibility criteria included being between 18 and 30 years of age or 60 years and older, having learned English before the age of 5, having normal or corrected-to-normal vision, not using hearing aids, and having no recent history of head trauma or neurological or psychological disorders. Four young adults and five older adults were excluded due to technical issues with either the fNIRS recording or the speech-in-noise task. The final sample consisted of 70 participants, including 36 young adults (age: *M* = 22.8 years, *SD* =3.1, range = 18-30; 22 females, 14 males) and 34 older adults (age: *M* = 74.2 years, SD =5.3, range = 63-83; 20 females, 14 males).

This sample size was informed by our previous study (Perron et al., 2025), which included 22 younger adults and 35 older adults. A simulation-based power analysis of the fitted linear mixed-effects model from that study, using the simr package in R with 1,000 simulations, estimated 86.2% power (95% CI [83.9%, 88.3%]) to detect the fixed effect of Age Group. Accordingly, we targeted a comparable sample size.

All participants were screened for cognitive status using the Montreal Cognitive Assessment Hearing Impairment version (MoCA-HI; Lin et al., 2017). A score of 25 or above is considered within the normal range. MoCA-HI scores were used to characterize the sample but were not applied as an exclusion criterion. MoCA scores did not differ significantly between age groups (Young adults: *M* = 28.32, *SD* = 1.77, range = 25 to 30; Older adults: *M* = 28.33, *SD* = 1.47, range = 23 to 30), *t*(68) = 0.03, *p* = 0.98.

In line with recent best practices in fNIRS research (Kwasa et al., 2023; Yücel et al., 2021), participants’ skin tone, hair type, and hair color were documented using self-report chart-based measures, while head circumference was measured directly. As these factors can influence fNIRS signal quality, the measures were collected for descriptive and methodological transparency only. Summary data for both groups are provided in the Supplementary Material 1.

### 2.2. Procedure

Participants were invited to attend a 2-hour session. First, they were asked to review the consent form. Then, they completed the MoCA test, which was administered by a trained research assistant in a quiet interview room. Next, they underwent hearing assessments and completed the main speech-in-noise task, during which their brain activity was recorded using fNIRS. These tests were conducted in a double-walled, soundproof booth. Participants were compensated for their time in the form of course credit or a small monetary incentive. Participants were allowed to take breaks as needed.

### 2.3. Hearing Assessments

All participants underwent a hearing screening procedure. This included an otoscopic examination to confirm the absence of obstructions or visible pathologies in the ear canal. Pure tone air conduction audiometry was conducted using a calibrated clinical audiometer (GSI 61, Grason-Stadler, USA). Pure-tone thresholds were measured in hearing levels (dB HL) at frequencies of 0.25, 0.5, 1, 2, 4, 6 and 8 kHz for each ear. The resulting audiograms are shown in Figure 1A. As expected, older adults exhibited significantly higher hearing thresholds than younger adults at all tested frequencies in both ears (all *ps*. < 0.05).

**Figure 1.**
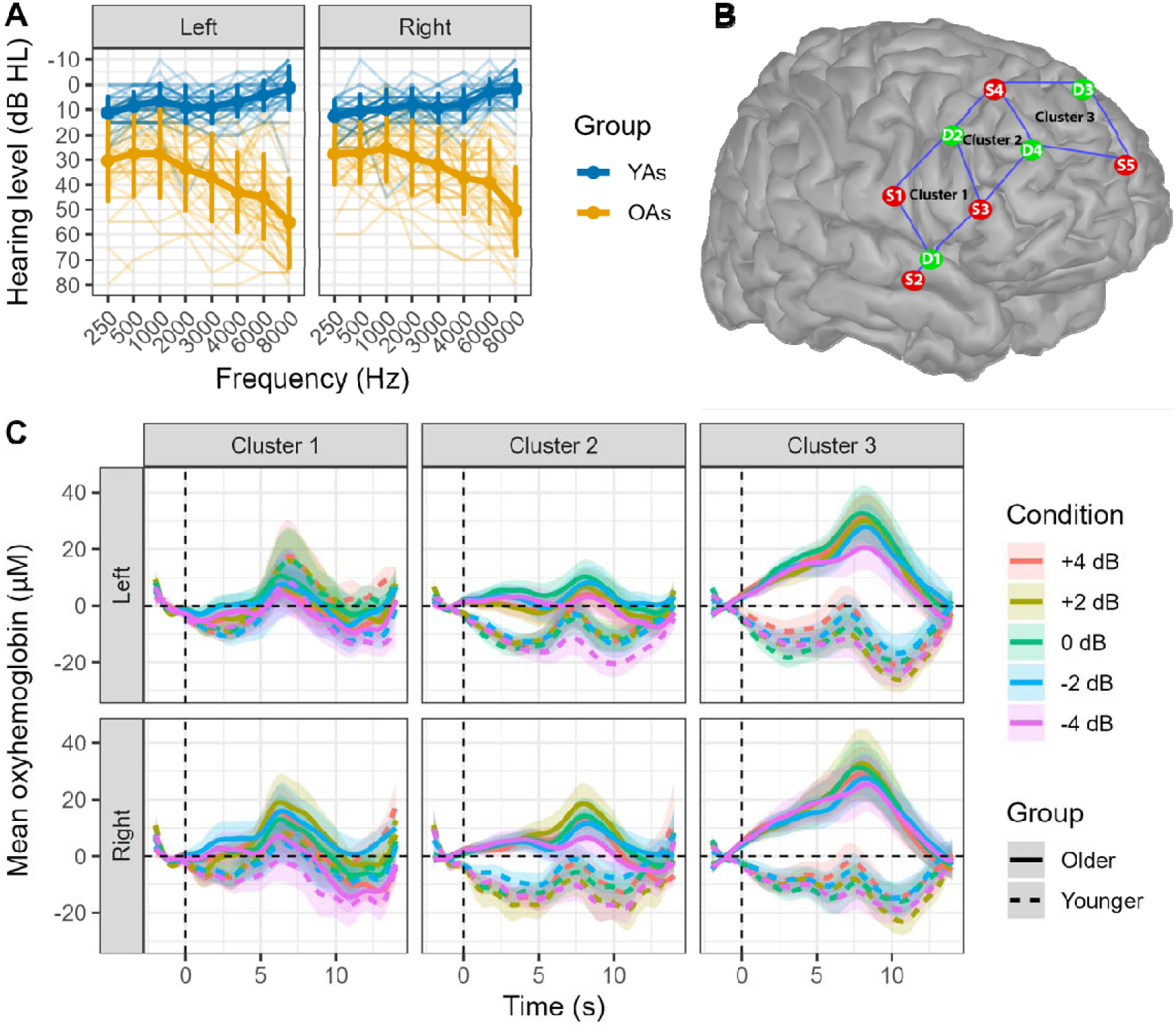
Audiometric assessment and fNIRS acquisition overview. (A) Pure-tone hearing thresholds (audiograms). (B) fNIRS probe configuration over the prefrontal cortex (right hemisphere only). Cluster 1 included channels S1-D1, S1-D2, and S3-D3. Cluster 2 included channels S3-D2, S3-D4, S4-D2, and S4-D4. Cluster 3 included channels S4-D3, S5-D3, and S5-D4. (C) Grand-average hemodynamic response functions (HRFs) time-locked to stimulus presentation for each cluster, signal-to-noise ratio condition and group. Shaded areas indicating the 95% confidence intervals.

A pure-tone average (PTA4) was calculated based on the average at four frequencies (0.5, 1, 2 and 4 kHz) (Humes, 2019; Stevens et al., 2013). Based on the PTA4 of the better ear (better ear PTA), older adults showed a distribution of hearing sensitivity spanning the normal (n = 10), mild (n = 18), moderate (n = 8), and moderately severe (n = 3) ranges. Hearing sensitivity among young adults was predominantly within the normal range, with one individual classified in the mild range.

### 2.4. Revised Speech-in-Noise (R-SPIN) Task

#### 2.4.1. Task Design

Speech-in-noise ability was assessed using an adapted version of the original R-SPIN task. Participants listened to sentences presented in multi-talker babble noise at the following SNRs: +4, +2, 0, −2 and −4 dB. A total of 120 trials were presented, with 24 trials per SNR condition. Trials from all SNR conditions were interleaved within each block in a mixed (event-related) design, and trial order was randomized across participants. The task was divided into six blocks of 20 trials to allow brief rest periods and minimize fatigue. The task was programmed in MATLAB 2022b (Natick, Massachusetts: The MathWorks Inc). Stimuli were presented through ER-3C insert earphones (Etymotic Research).

Sentence materials varied in contextual predictability. In half of the trials, the final word was highly constrained by the preceding context, whereas in the remaining trials it was not. This manipulation was included to minimize ceiling and floor effects, promote sustained attention across the entire sentence, and reduce the likelihood of strategic listening. When context is highly constraining, listeners may anticipate the final word before it is presented. In contrast, sentences with little contextual support may reduce engagement if they provide limited predictive information. Context predictability was not treated as a variable of interest, consistent with our previous study in which no effect of context was observed in our regions of interest (Perron et al., 2025).

Each trial began with 1 s of babble noise, followed by the target sentence, and ended with an additional 2 s of babble noise. The babble included 50 ms linear onset and offset ramps, Participants were instructed to wait until the stimulus had ended before repeating aloud the final word of the sentence. If they were unsure, they were encouraged to provide their best guess or to respond “pass.” After their verbal response, participants pressed the space bar to proceed to a listening effort rating screen. They rated the effort required to understand the final word using the 7-point scale developed by Johnson et al. (2015), where 1 corresponded to “No effort” and 7 to “Extreme effort.” Participants were instructed to rate their perceived listening effort rather than the accuracy of their response. Following the effort rating, the next trial began after a randomly jittered intertrial interval ranging from 8 to 12 s.

#### 2.4.2. Individualized Stimulus Calibration

Stimulus presentation levels were individually calibrated to reduce differences in audibility across participants. Specifically, presentation levels were normalized to each participant’s behavioural detection threshold for the multi-speaker background noise used in the main task (i.e., the babble threshold; Russo and Pichora-Fuller, 2008). This approach is analogous to standard pure-tone audiometry, except that thresholds were established for broadband background noise rather than individual frequencies.

Babble thresholds were measured separately for each ear. The babble noise was initially presented at 50 dB HL and adjusted in 10 dB steps until detected. The level was then decreased by 10 dB and increased in 5 dB steps until detection was reestablished. Threshold was defined as the presentation level corresponding to approximately 50% detection performance (2 detections in 4 presentations, or 3 in 6 if the participant reported uncertainty).

During the main task, target speech was presented at a sensation level of 50 dB above each participant’s babble threshold. The background noise level was then varied relative to this fixed speech level to produce the desired SNRs. This calibration minimized variability attributable to baseline audibility while maintaining equivalent relative differences in listening difficulty across SNR conditions. Importantly, the procedure was not designed to equate behavioural performance across participants. All participants completed the same SNR conditions, preserving individual differences in speech percep tion for subsequent behavioural and brain-behaviour analyses.

### 2.5. fNIRS Data Acquisition

The fNIRS data were acquired using a continuous wave optical system (Dual Brite MKIII, Artinis Medical Systems, Netherlands) operating at 756 nm and 842 nm, with a sampling rate of 100 Hz (which was down sampled to 10 Hz). Two probes with identical montages were used. Each probe consisted of ten light sources and eight detectors arranged with a 3 cm inter-optode distance. One probe was positioned over the bilateral prefrontal cortex, with channels evenly distributed in a 2 × 10 channel configuration (Figure 1B). A second probe was positioned over the parietal and temporal cortices but was not analyzed further due to excessive noise in the young adult group. Each probe also included two short-separation channels (one in each hemisphere) with a 1 cm inter-optode distance to capture superficial hemodynamic fluctuations from the scalp. Probe placement was guided by the fNIRS Optodes’s location Decider (fOLD) toolbox (Zimeo Morais et al., 2018) using the Jülich Atlas.

Participants wore small (N = 6; 53-55 cm), medium (N = 43; 55-57 cm), or large (N = 21; 57-59 cm) caps. Cap placement was standardized using the international 10-20 system. The Cz position was determined by measuring the distance between the nasion and the inion and marking its midpoint. The distance between the left and right preauricular points were used to ensure lateral symmetry. The Cz marker on the fNIRS cap was then precisely aligned with the participant’s Cz.

Optode positions and anatomical landmarks (nasion, inion, and bilateral preauricular points) were digitized for 62 participants (31 younger adults and 31 older adults) using a 3-D digitizer (Patriot, Polhemus, Colchester, VT) within Brainstorm (Tadel et al., 2011). For each participant, additional head shape points were collected and used to warp a standard MNI template to individual anatomy. Source-detector positions were then used to estimate participant-specific channel locations on the cortical surface. Group-averaged MNI coordinates for each optode and channel are reported in Supplementary Material 2.

### 2.6. fNIRS Data Preprocessing

Data preprocessing and analysis followed a pipeline similar to our prior work (Perron et al., 2025), and were carried out in Homer3 (v 1.87.0; Huppert et al., 2009). Channels with low SNR were excluded using the hmrR_PruneChannels function (dRange = [−1e2, 1e7]; SNRthresh = 2). Raw intensity signals from the remaining channels were converted to optical density using hmrR_Intensity2OD. Motion artifacts were corrected using a wavelet-based approach (hmrR_MotionCorrectWavelet; iqr = 1.5). To attenuate high-frequency physiological noise, optical density data were low-pass filtered at 0.5 Hz using hmrR_BandpassFilt.

Variations in oxygenated (HbO) and deoxygenated (HbR) hemoglobin concentrations were computed using the Modified Beer-Lambert law (hmrR_OD2Conc; ppf = 1), with values expressed as concentration change multiplied by mean optical path length. Hemodynamic responses were estimated using a general linear model over a −2 to 14 s time window relative to stimulus onset. The model was solved using a least-squares approach with a set of temporally shifted Gaussian basis functions (hmrR_GLM; glmSolveMethod = 1, idxBasis = 1). Slow signal drifts were modeled using a third-order polynomial, and baseline correction was incorporated directly into the general linear model using the −2 to 0 s pre-stimulus interval. To reduce the influence of systemic physiological activity, short-separation regression was applied by selecting, for each long-separation channel, the short-separation channel exhibiting the highest correlation (flagSSmethod = 1). In rare cases where no short-separation channels remained, short-separation regression could not be performed, and the affected long-separation channels were retained without correction. The resulting beta coefficients were combined with their associated basis functions to reconstruct the full hemodynamic response function time course.

Following preprocessing, the long-separation channels were grouped into three prefrontal clusters, based on the geometry of the probes and their correspondence with the underlying cortical regions (Supplementary Material 2). The channel assignments are described in Figure 1B. The analysis focused *a priori* on Cluster 2, which corresponded to the DLPFC and comprised channels S3-D2, S3-D4, S4-D2 and S4-D4. This cluster matched the region of interest examined in our previous study (Perron et al., 2025). We defined additional exploratory regions of interest. Cluster 1 corresponded to the VLPFC and included channels S1-D1, S1-D2 and S3-D3. Cluster 3 corresponded to the DMPFC and included channels S4-D3, S5-D3 and S5-D4. The same clustering scheme was applied to the left hemisphere.

Prior to cluster averaging, outlier values exceeding 2.5 times the interquartile range above or below the mean were removed for each channel independently to limit the influence of extreme responses. HbO and HbR signals were then averaged across the remaining channels within each cluster for each participant and condition. Participants were included in a given cluster if at least one long-separation channel remained within that cluster.

Figure 1C shows the HbO time courses for each SNR, age group, and cluster, while Supplementary Material 3 presents the corresponding HbR time courses. To facilitate replication of our previous study, the primary analyses focused on HbO. This approach was also motivated by evidence that HbO exhibits greater test-retest reproducibility than HbR (Plichta et al., 2006) and corresponds more closely to the fMRI BOLD response (Cui et al., 2011). As shown in the Supplementary Material 3, HbR responses generally showed the expected inverse pattern relative to HbO, although they were smaller in amplitude and visibly noisier. Parallel analyses of the HbR responses are reported in the Supplementary Material 4.

HbO responses were averaged over a post-stimulus window ranging from 5.5 to 11 s. This window was selected following a visual inspection of the overall HbO time curve (Figure 1C) in order to capture the peak of the task-induced hemodynamic response. Compared to our previous study, the peak occurred approximately 2 s earlier, resulting in a shift forward in the analysis window (5.5–11 s versus 7.5–13 s). This difference is likely related to differences between the tasks in the two studies, notably the change from a block-design experimental protocol to a mixed/event-related design, as well as to the range and distribution of SNR conditions.

### 2.7. Statistical Analyses

Behavioural and fNIRS data analyses were performed in R using RStudio (RStudio Team, 2022). The distribution of each dependent variable was visually inspected using histograms and Q-Q plots to assess approximate normality and identify potential deviations from model assumptions. No substantial violations were observed that warranted transformation of the data. Outliers were identified using a 2.5× interquartile range criterion applied separately within each factor of interest included in the corresponding analyses.

All main analyses were conducted using mixed-effects models fitted with the *lmer* function from the *lme4* package. In all models, SNR was coded as an ordered numeric predictor. The linear SNR term was mean-centred, and the quadratic SNR term was calculated from the centred linear term and subsequently mean-centred. Participant was included as a random intercept. To identify the most parsimonious model, we used an information-theory-based model selection approach, based on the corrected Akaike information criterion (AICc) implemented in the *MuMIn* package (*dredge* function). The candidate models were fitted using maximum likelihood estimation. The models were ranked using AICc, which balances model fit and complexity while applying an additional correction for finite sample sizes (Burnham and Anderson, 2002).

For each analysis, two candidate model sets were evaluated. The first considered models containing only the linear SNR term and its interactions with the relevant grouping factors. The second additionally included the quadratic SNR term and its interactions, while enforcing polynomial hierarchy such that the linear SNR term was retained whenever the quadratic term was present. The highest-ranked model from each candidate set was then compared, and the model with the lower AICc and higher Akaike weight was selected for subsequent inference.

For the selected models, fixed effects were evaluated using Satterthwaite’s approximation for denominator degrees of freedom. Significant effects were followed up with estimated marginal means and pairwise contrasts using the *emmeans* package. Fixed-effect estimates are reported as standardized regression coefficients (β) with their standard errors and 95% confidence intervals. Variance inflation factors (VIFs) were examined for all selected models and indicated negligible multicollinearity (all adjusted VIFs ≤ 1.52).

#### 2.7.1. Behavioural Analyses

Separate analyses were conducted for accuracy and self-reported listening effort. For each outcome, two candidate model sets were evaluated. The global linear model included the fixed effects of Group, linear SNR, and their interaction (Outcome ∼ Group × linear SNR + (1 | Participant)), whereas the global quadratic model additionally included the quadratic SNR term and its interactions (Outcome ∼ Group × (linear SNR + quadratic SNR) + (1 | Participant)).

#### 2.7.2. fNIRS Analyses

For the fNIRS data, three complementary analytic approaches were used, with separate LMMs fitted for each cluster.

First, mean HbO responses were analysed to characterize age-related differences in task-related cortical activation across SNR conditions. The global linear model included the fixed effects of Group, Hemisphere, linear SNR, and all interactions (HbO ∼ Group × Hemisphere × linear SNR + (1 | Participant)), whereas the global quadratic model additionally included the quadratic SNR term and its interactions (HbO ∼ Group × Hemisphere × (linear SNR + quadratic SNR) + (1 | Participant)).

Second, to examine interindividual variability within the older adult group, these analyses were repeated after replacing the two-level Group factor with a three-level Group factor comprising younger adults, low-performing older adults, and high-performing older adults. The older adult group was subdivided using a median split of overall accuracy. To characterize the subgroups, participant characteristics were compared using Welch’s independent-samples t-tests. The low-performing and high-performing older adult subgroups did not differ significantly in age (low-performing: *M* = 75.7, *SD* = 4.6 years; high-performing: *M* = 73.0 ± 5.1 years; *t*(31.63) = 1.63, *p* = .113), Better ear PTA (low-performing: *M* = 33.9, *SD* = 11.8 dB HL; high-performing: *M* = 27.0, *SD* = 11.2 dB HL; *t*(31.90) = 1.76, *p* = .087), or MoCA score (low-performing: *M* = 27.9, *SD* = 1.7; high-performing: *M* = 28.7, *SD* = 1.8; *t*(31.93) = −1.27, *p* = .213).

Third, to determine whether neural responses differed as a function of behavioural outcome, mean HbO responses were averaged separately across correct and incorrect trials, and the primary fNIRS models were extended to include Trial Type as an additional fixed effect in both global linear and quadratic models.

#### 2.7.3. Brain-Behaviour Relationships

Brain-behaviour relationships were examined using partial correlations controlling for age, better ear PTA and MoCA to assess the relationships between HbO responses and behavioural measures. Partial Pearson correlations were computed for accuracy, which was treated as a continuous variable, whereas partial Spearman correlations were computed for listening effort because effort ratings were measured on an ordinal scale.

Additional Pearson correlations were performed to examine the relationships between HbO responses and age, better ear PTA, and MoCA scores. All analyses were performed separately within each Age Group, Hemisphere, and SNR condition.

False discovery rate (FDR) correction was applied within each Age Group and Hemisphere to correct for multiple comparisons.

## 3. Results

### 3.1. Behavioural Performance

#### 3.1.1. Accuracy

For accuracy, the best-supported linear model included the fixed effects of Group and the linear SNR term (Accuracy ∼ Group + linear SNR + (1 | Participant)), whereas the best-supported hierarchical quadratic model additionally retained the quadratic SNR term (Accuracy ∼ Group + linear SNR + quadratic SNR + (1 | Participant)). Comparison of the two models using a likelihood-ratio test indicated that adding the quadratic SNR term significantly improved model fit (χ*²*(1) = 83.51, *p* < .001). Consistent with this result, the hierarchical quadratic model also showed substantially greater support based on information-theoretic model selection (AICc = 2640.6) than the corresponding linear model (AICc = 2722.0; ΔAICc = 81.4; Akaike weight = 1.00).

Results are shown in Figure 2A. The selected model revealed significant linear (β = −11.35, *SE* = 0.33, *95% CI* [−12.01, −10.69], *t*(344) = −33.98, *p* < .001) and quadratic (β = −2.78, *SE* = 0.28, *95% CI* [−3.34, −2.23], *t*(344) = −9.86, *p* < .001) effects of SNR, indicating that accuracy declined as listening conditions became more difficult, with the rate of decline becoming progressively steeper at poorer SNRs. A significant main effect of Group was also observed (β = −6.29, *SE* = 2.07, *95% CI* [−10.36, −2.21], *t*(344) = −3.04, *p* = .003), with younger adults demonstrating higher overall accuracy (*M* = 64.0%) than older adults (*M* = 57.7%).

**Figure 2.**
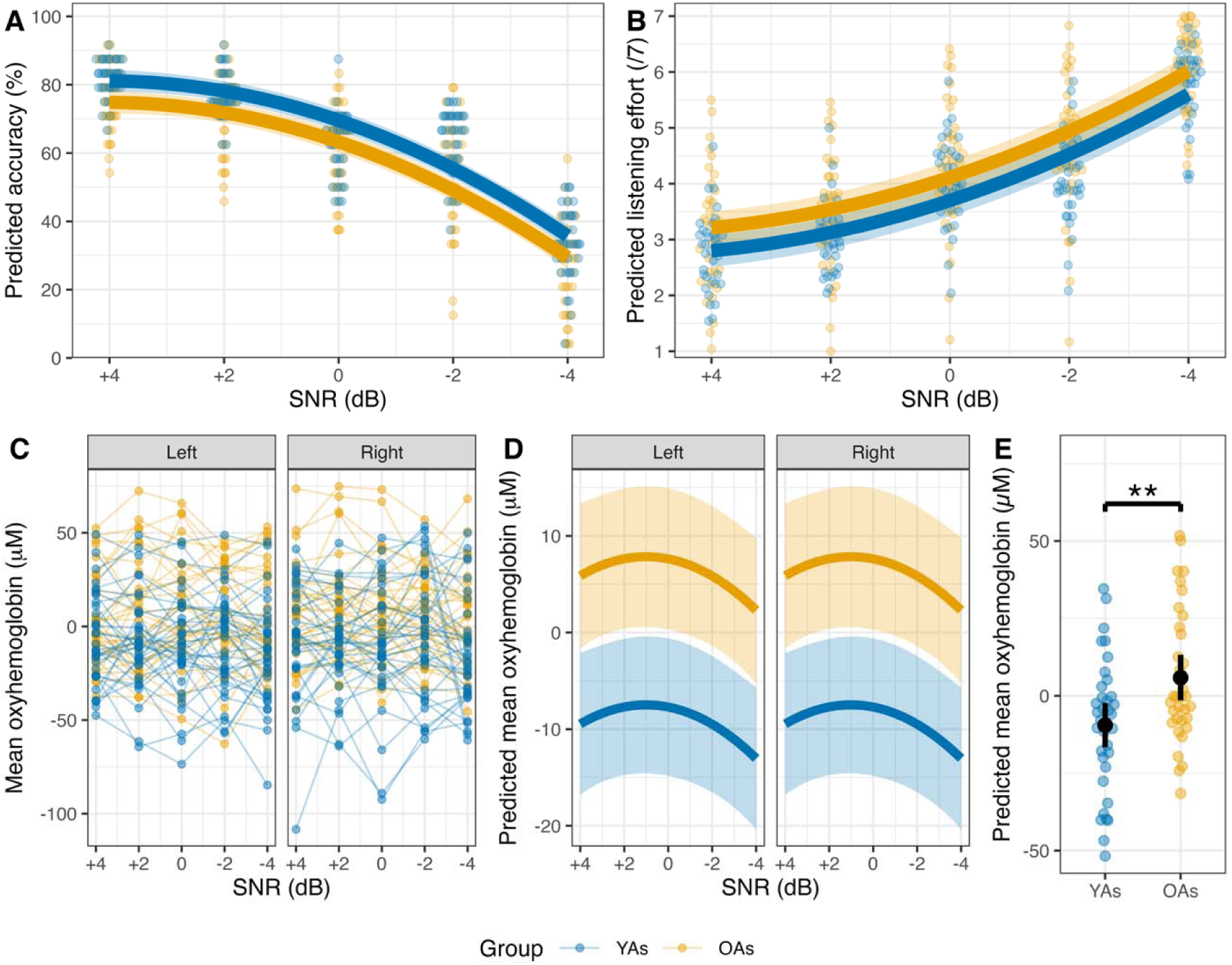
Behavioural and neural responses for younger adults (YAs) and older adults (OAs) across signal-to-noise ratio (SNR) conditions. (A) Accuracy. (B) Self-reported listening effort. (C) Individual dorsolateral prefrontal cortex (DLPFC) oxyhemoglobin (HbO) responses across SNR conditions, with lines connecting observations from the same participant. (D) Group-level predicted DLPFC HbO responses from the final mixed-effects model. Solid lines represent predicted values, and shaded areas indicate 95% confidence intervals. (E) Estimated marginal means for DLPFC HbO by group, with points representing individual participant-level mean HbO values and error bars representing 95% confidence intervals. **p < 0.01,

#### 3.1.2. Self-Reported Listening Effort

For self-reported listening effort, the same best-supported model structures as for accuracy were compared. Comparison of these models indicated that the hierarchical quadratic model provided a significantly better fit to the data than the best linear model (χ*²*(1) = 66.56, *p* < .001). The hierarchical quadratic model also received substantially greater support based on information-theoretic model selection (AICc = 646.9 vs. 711.4; ΔAICc = 64.5; Akaike weight = 1.00).

Results are shown in Figure 2B. The selected model revealed significant linear (β = 0.70, *SE* = 0.02, *95% CI* [0.67, 0.73], *t* = 40.90, *p* < .001) and quadratic (β = 0.13, *SE* = 0.01, *95% CI* [0.10, 0.15], *t* = 8.66, *p* < .001) effects of SNR, indicating that self-reported listening effort increased as listening conditions became more difficult, with the increase becoming progressively steeper at poorer SNRs. A significant main effect of Group was also observed (β = 0.41, *SE* = 0.20, *95% CI* [0.02, 0.81], *t* = 2.05, *p* = .041), with older adults (*M* = 4.36) reporting greater listening effort than younger adults (*M* = 3.95).

### 3.2. Effect of SNR and Age on DLPFC Oxygenation

For DLPFC HbO, the best-supported linear model included the fixed effects of Group and the linear SNR term (HbO ∼ Group + linear SNR + (1 | Participant)), whereas the best-supported hierarchical quadratic model also included the quadratic SNR term (HbO ∼ Group + linear SNR + quadratic SNR + (1 | Participant)). Comparison of these models indicated that the hierarchical quadratic model provided a significantly better fit to the data than the best linear model (χ*²*(1) = 5.26, p = .022). The hierarchical quadratic model also received greater support based on information-theoretic model selection (AICc = 6090.6 vs. 6093.8; ΔAICc = 3.23; Akaike weight = 0.834).

Individual HbO responses across SNR conditions are shown in Figure 2C, and the corresponding model-predicted trajectories are shown in Figure 2D. The fitted trajectory indicated that HbO responses increased from favourable to intermediate listening conditions before declining under the most challenging listening conditions. This pattern was captured by significant linear (β = −0.90, *SE* = 0.46, *95% CI* [−1.80, −0.01], *t* = −1.98, *p* = .048) and quadratic (β = −0.88, *SE* = 0.38, *95% CI* [−1.64, −0.13], *t* = −2.30, *p* = .022) terms. A significant main effect of Group was also observed (β = 15.33, *SE* = 5.09, *95% CI* [5.34, 25.32], *t* = 3.01, *p* = .003), with older adults exhibiting greater overall HbO responses (*M* = 5.85 µM) than younger adults (*M* = −9.48 µM) (Figure 2E). Exploratory pairwise comparisons further confirmed significantly greater HbO responses in older adults than in young adults across all five SNR levels (all *p*s < .018).

An exploratory analysis including better ear PTA and MoCA scores as covariates yielded the same model selection results.

### 3.3. High-Performing vs. Low-Performing Older Adults

Additional analyses were conducted using a three-level group factor comprising younger adults, high-performing older adults, and low-performing older adults. Older adults were divided into high- and low-performing subgroups using a median split on overall accuracy (median = 59.9%). The behavioural performance of the three groups is shown in Figure 3A-B. As illustrated, the high-performing older adults exhibited behavioural performance comparable to that of younger adults across listening conditions, whereas the low-performing older adults showed lower accuracy and greater listening effort across all SNRs.

**Figure 3.**
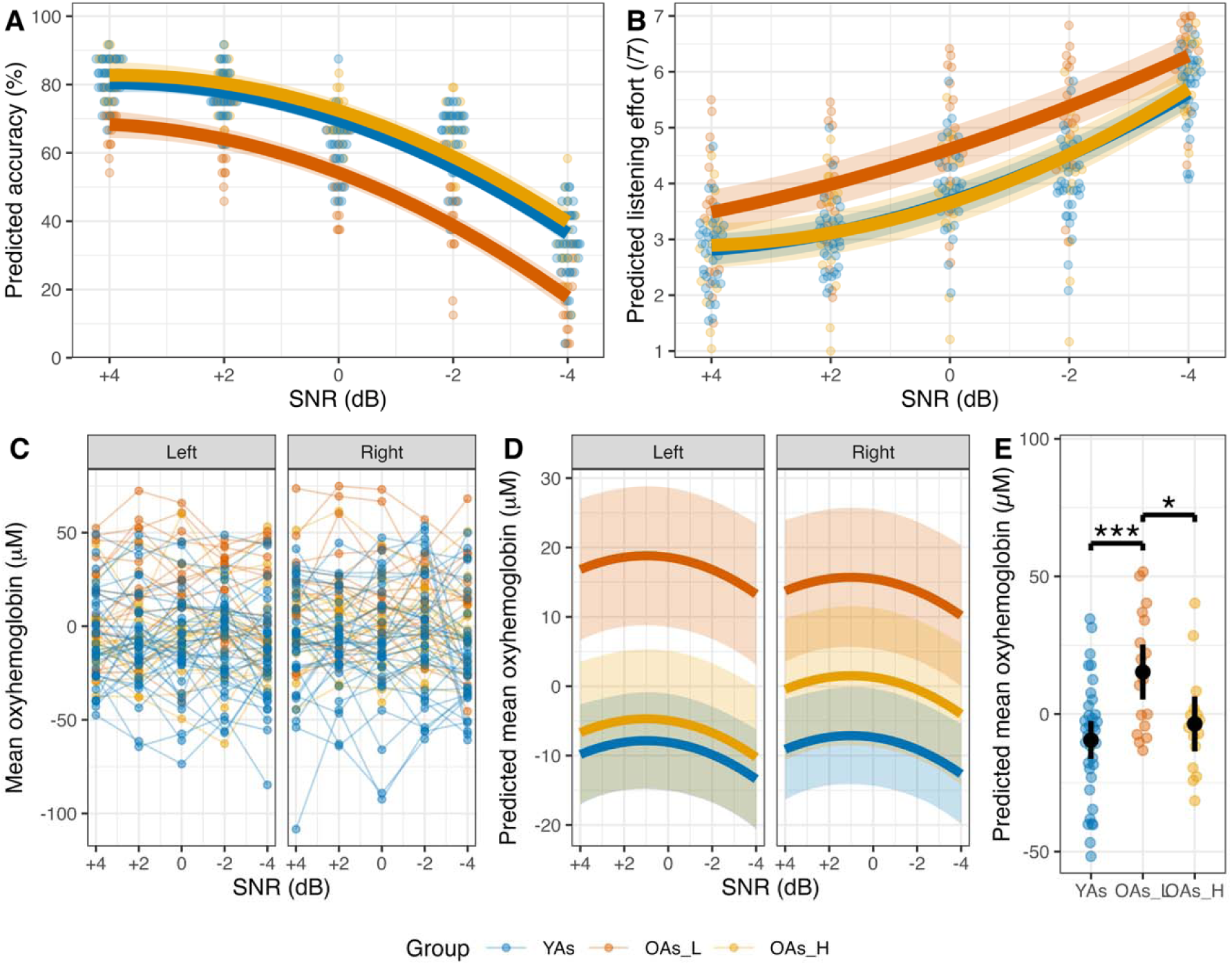
Behavioural and neural responses for younger adults (YAs), low-performing older adults (OAs_L), and high-performing older adults (OAs_H) across signal-to-noise ratio (SNR) conditions. (A) Accuracy. (B) Self-reported listening effort. (C) Individual dorsolateral prefrontal cortex (DLPFC) oxyhemoglobin (HbO) responses across SNR conditions, with lines connecting observations from the same participant. (D) Group-level predicted DLPFC HbO responses from the final mixed-effects model. Solid lines represent predicted values, and shaded areas indicate 95% confidence intervals. (E) Estimated marginal means for DLPFC HbO by group, with points representing individual participant-level mean HbO values and error bars representing 95% confidence intervals. *p < 0.05, ***p < 0.001.

For HbO, the best-supported linear model included the fixed effects of Hemisphere, Group, and the linear SNR term, together with the Group × Hemisphere interaction (HbO ∼ Hemisphere + Group + linear SNR + Group × Hemisphere + (1 | Participant)). The best-supported hierarchical quadratic model additionally retained the quadratic SNR term (HbO ∼ Hemisphere + Group + linear SNR + quadratic SNR + Group × Hemisphere + (1 | Participant)). Comparison of these models indicated that the hierarchical quadratic model provided a significantly better fit to the data than the best linear model (χ*²*(1) = 5.32, *p* = .021). The hierarchical quadratic model also received greater support based on information-theoretic model selection (AICc = 6084.4 vs. 6087.7; ΔAICc = 3.27; Akaike weight = 0.837).

Individual HbO responses across SNR conditions are shown in Figure 3C, and the corresponding model-predicted trajectories are shown in Figure 3D. Consistent with the previous analysis, the fitted HbO trajectory exhibited a nonlinear relationship with SNR, with responses increasing from favourable to intermediate listening conditions before declining under the most challenging listening conditions. This pattern was captured by significant linear (β = −0.90, *SE* = 0.45, *95% CI* [−1.79, −0.01], *p* = .047) and quadratic (β = −0.88, *SE* = 0.38, *95% CI* [−1.63, −0.13], *p* = .021) terms.

A significant main effect of Group was observed for the low-performing older adult subgroup (β = 22.83, *SE* = 6.16, *95% CI* [10.73, 34.92], *p* < .001), whereas the contrast between high-performing older adults and younger adults was not significant (β = 8.64, *SE* = 6.16, *95% CI* [−3.46, 20.73], *p* = .161). Mean HbO responses were highest among low-performing older adults (*M* = 15.27 µM), followed by high-performing older adults (*M* = −3.58 µM) and younger adults (*M* = −9.47 µM) (Figure 3E).

The main effect of Hemisphere was not significant (β = 0.74, *SE* = 1.80, *95% CI* [−2.79, 4.28], *p* = .681). However, the omnibus Group × Hemisphere interaction was significant, *F*(2, 608.50) = 3.28, *p* = .038. The corresponding interaction coefficients were not individually significant for either low-performing older adults (β = −3.84, *SE* = 3.14, *95% CI* [−10.01, 2.34], *p* = .223) or high-performing older adults (β = 5.48, *SE* = 3.14, *95% CI* [−0.69, 11.66], *p* = .082).

Exploratory post hoc comparisons revealed that, in the left hemisphere, low-performing older adults exhibited significantly greater HbO responses than both younger adults (*p* = .001) and high-performing older adults (*p* = .022), whereas younger adults and high-performing older adults did not differ (*p* = .996). In the right hemisphere, low-performing older adults also exhibited greater HbO responses than younger adults (*p* = .006), but did not differ from high-performing older adults (*p* = .387). High-performing older adults and younger adults did not differ (*p* = .743). No significant hemispheric differences were observed within any group (all *ps* ≥ .157).

An exploratory analysis including better ear PTA and MoCA scores as covariates yielded the same model selection results.

### 3.4. Correct vs. Incorrect Responses

When Trial Type was added to the candidate model set, it was not retained in either the best-supported linear or hierarchical quadratic model. The selected model had the same fixed-effects structure as the primary fNIRS model and again indicated a nonlinear relationship between SNR and HbO responses, together with an overall Age Group difference.

### 3.5. Brain-Behaviour Relationships

#### 3.5.1. Correlations between Performance and DLPFC Oxygenation

We examined the relationships between accuracy, self-reported listening effort, and HbO responses in the DLPFC using partial correlations controlling for better ear PTA and MoCA. Results are shown in Figure 4A.

**Figure 4.**
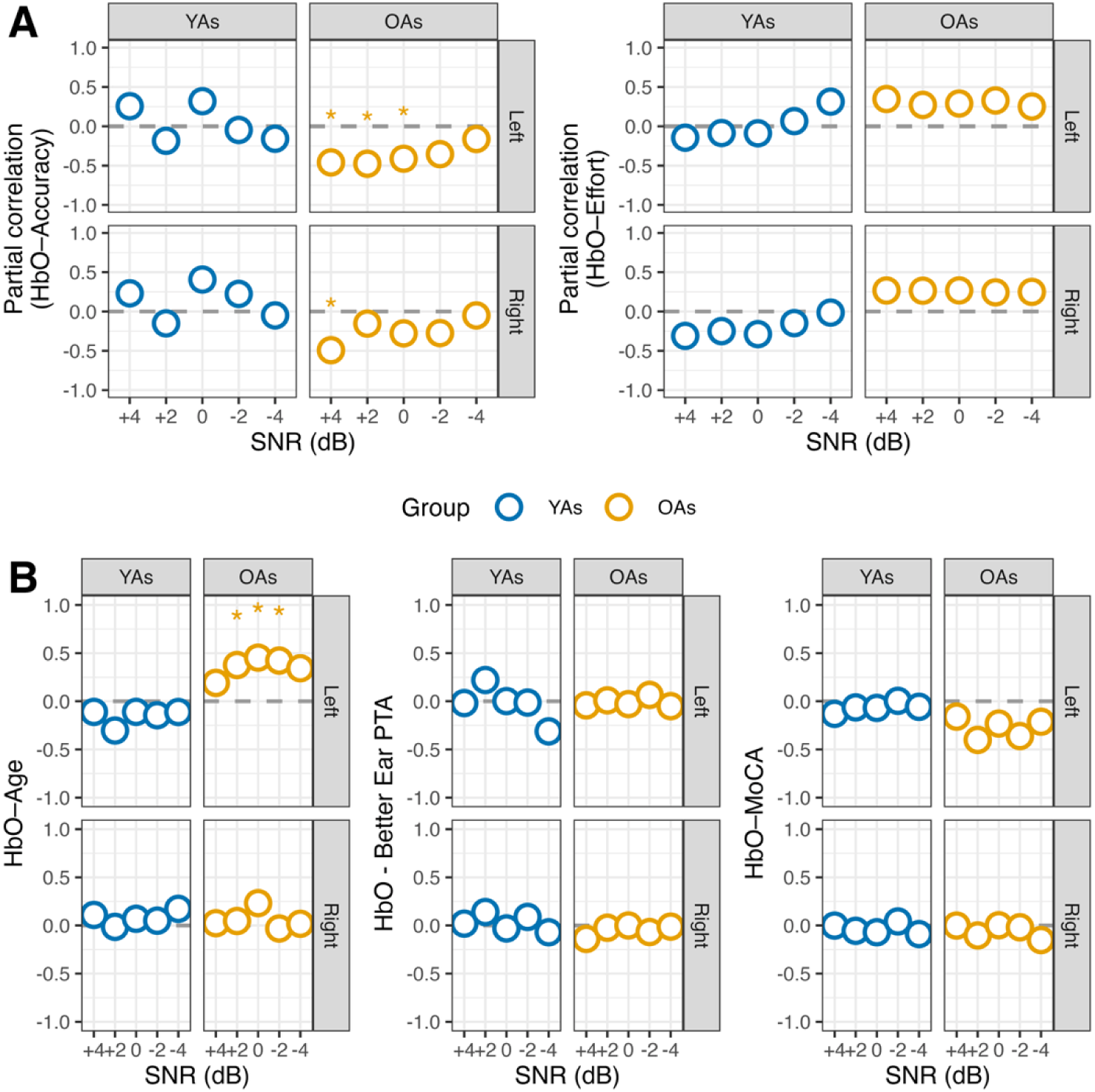
Brain-behaviour and demographic relationships. (A) Partial correlation coefficients between Dorsolateral prefrontal cortex (DLPFC) oxyhemoglobin (HbO) responses and behavioural measures (accuracy and self-reported listening effort) within each SNR condition and hemisphere, controlling for age, better ear pure-tone average (PTA) and cognition (MoCA scores). (B) Pearson correlation coefficients between DLPFC HbO responses and demographic measures (age, Better ear PTA, and MoCA scores). Asterisks indicate correlations that remained significant after false discovery rate correction. * p < 0.05, ** p < 0.01, *** p < 0.001.

In older adults, we observed significant negative associations between left DLPFC HbO responses and accuracy at +4 dB (*r* = −0.46, *p_FDR* = 0.023), +2 dB (*r* = −0.48, *FDR* = 0.023), and 0 dB SNR (*r* = −0.41, *FDR* = 0.035). The association at −2 dB SNR was similar in direction but did not survive FDR correction (*r* = −0.36, *FDR* = 0.063), and no association was observed at −4 dB SNR (*r* = −0.16, *FDR* = 0.385). Thus, greater left DLPFC oxygenation was generally associated with poorer speech-in-noise performance.

In the right DLPFC, older adults showed a significant negative correlation between HbO responses and accuracy at +4 dB SNR (*r* = −0.49, *FDR* = 0.025), whereas correlations at the remaining SNRs were not significant (all *FDR* ≥ 0.225).

In contrast, no significant correlations between HbO responses and accuracy were observed in young adults in either hemisphere across any SNR condition (all |*r*| < 0.41, all *p_FDR* > 0.10).

For listening effort, no significant correlations with HbO responses were found in either age group across any SNR condition or hemisphere (all |*r*| < 0.35, all *p_FDR* > 0.165).

#### 3.5.2. Correlations between Demographic Factors and DLPFC Oxygenation

Here, we examined whether DLPFC HbO responses were associated with individual difference factors including age, better ear PTA, and MoCA scores. Results are shown in Figure 4B.

In older adults, age was significantly and positively correlated with left DLPFC HbO responses at +2 dB (*r* = 0.38, *p_FDR* = 0.047), 0 dB (*r* = 0.45, *p_FDR* = 0.032), and 2 dB SNR (*r* = 0.42, *p_FDR* = 0.032), but not at +4 dB (*r* = 0.19, *p_FDR* = 0.280) or 4 dB SNR (*r* = 0.34, *p_FDR* = 0.059). Thus, greater age within the older adult group was associated with greater left DLPFC oxygenation. No significant correlations with age were observed in the right hemisphere in older adults or in younger adults in either hemisphere (all *p_FDR* ≥ 0.359).

Better ear PTA was not significantly correlated with HbO responses in either Age Group at any SNR or Hemisphere (all |*r*| ≤ 0.31, all *p_FDR* ≥ 0.319).

Similarly, MoCA scores were not significantly correlated with HbO responses in either age group after FDR correction (all *p_FDR* ≥ 0.089). In older adults, negative correlations were observed in the left hemisphere at +2 dB (*r* = −0.40, *p* = 0.019) and −2 dB SNR (r = −0.36, *p* = 0.036), but these effects did not survive the FDR correction. No other associations were observed (all |r| ≤ 0.23, all *p_FDR* ≥ 0.285).

### 3.6. Exploratory analyses: VLPFC and DMPFC Oxygenation

Exploratory analyses were conducted for the VLPFC (Cluster 1) and DMPFC (Cluster 3). Details results are provided and illustrated in Supplementary Material 5.

In the VLPFC, the hierarchical quadratic model provided a better fit to the data, indicating a non-linear relationship between HbO responses and SNR similar to that observed in the DLPFC. However, unlike the DLPFC, no overall differences were observed between younger and older adults. The only subgroup-specific effect was a Hemisphere × Group interaction, indicating that low-performing older adults exhibited relatively greater right-than left-hemisphere HbO responses. VLPFC HbO responses were also significantly related to age, better ear PTA, or MoCA scores.

In the DMPFC, the linear model provided a better fit to the data, revealing a decrease in HbO responses with increasing listening difficulty. Older adults exhibited greater overall HbO responses than younger adults, and this difference was evident in both the high- and low-performing older adult subgroups. DMPFC HbO responses were also not related to age, better ear PTA, or MoCA scores.

## 4. Discussion

This study investigated whether age-related prefrontal activity during speech-in-noise perception reflects compensatory recruitment, neural inefficiency, or disengagement within a non-linear framework of listening difficulty. Using fNIRS, we measured HbO responses in three functionally distinct prefrontal subregions, the DLPFC, VLPFC, and DMPFC, across five SNR conditions in young and older adults. In the DLPFC, HbO responses followed an inverted U-shaped function of listening difficulty, with older adults showing significantly greater responses across all SNR conditions. Critically, greater DLPFC recruitment in older adults was associated with poorer accuracy across the ascending and peak portions of the response function. These findings suggest that age-related DLPFC recruitment is more consistent with a neural inefficiency account than with compensatory recruitment and extend our previous work by demonstrating that this relationship is embedded within a non-linear, demand-dependent profile of listening difficulty. The VLPFC and DMPFC showed distinct patterns of modulation by age and SNR, suggesting partially dissociable functional roles. However, activity in these regions did not significantly predict behavioural performance in either age group, indicating that the strongest evidence for neural inefficiency was observed in the DLPFC. Taken together, these findings suggest that the DLPFC response may comprise two distinguishable features: its overall trajectory reflects changes in recruitment across listening demand, whereas its magnitude reflects age-related differences in neural recruitment associated with performance.

### 4.1. Inverted U-Shaped Relationship in the DLPFC

The results support our first hypothesis that DLPFC responses would follow an inverted U-shaped trajectory across SNR conditions. In all models, the hierarchical model provided a better fit to the data than the corresponding linear model, supporting the investigation of non-linear neural responses. We observed that HbO responses increased at intermediate levels of difficulty before declining under the most adverse SNR conditions (peaking at approximately 0 dB SNR). This non-linear profile aligns with converging evidence showing inverted U-shaped relationships between task demand and other indices of listening effort, including dual-task reaction time (Wu et al., 2016), frontal and parietal alpha power (Paul et al., 2021; Ryan et al., 2022), and pupil dilation (Zekveld and Kramer, 2014). To the best of our knowledge, this is the first evidence of such a pattern in terms of hemodynamic response in the DLPFC. Although the magnitude of the quadratic term was modest, this result is consistent with those of the studies mentioned above, in which the nonlinear effects were also found to be relatively weak. Using fNIRS, Lawrence et al. (2018), in young adults, similarly reported significant quadratic effects in bilateral temporal and parietal regions, and only a non-significant inverted U-shaped pattern in the left inferior frontal cortex.

Traditionally, this non-linear pattern has been interpreted as reflecting the progressive and beneficial recruitment of cognitive resources as task demands increase, followed by a decline once available capacity is exceeded. This interpretation has generally relied on the shape of the response alone, with little consideration of how each point along the curve relates to performance. The present study addressed this gap. Under a compensatory account, greater activation along the ascending portion of the curve should be accompanied by better or maintain behavioural performance, reflecting the successful mobilization of cognitive resources. Instead, we found that greater DLPFC activation was associated with poorer accuracy throughout the ascending and peak portions of the demand-response function, providing little evidence that increasing activation reflects successful compensatory recruitment.

The CRUNCH model similarly predicts that older adults recruit additional neural resources at lower levels of demand but reach capacity limits earlier than younger adults because of reduced resources (Reuter-Lorenz and Cappell, 2008). Our findings also provide limited support for this account. The quadratic relationship between DLPFC activation and listening demand was preserved across age groups and older adult subgroups, indicating that aging did not alter the shape of the demand-response function. Rather, age differences were primarily reflected in the magnitude of the response, with older adults showing greater overall activation across listening conditions. This pattern is consistent with a recent functional MRI study testing the CRUNCH model in a visuospatial working memory task across four levels of difficulty (Jamadar, 2020). Jamadar found that age-related differences in activation generally increased with task difficulty. More specifically, several frontoparietal regions, including the superior and middle frontal gyri, precentral gyrus, and superior parietal lobule, exhibited inverted U-shaped response profiles, with activation increasing from low to intermediate loads before declining at the highest load. Importantly, older adults did not show an earlier downturn in activation compared to younger adults. Rather, they exhibited greater activation at the highest load levels. This finding mirrors the current results. It is unlikely that the absence of a CRUNCH-like pattern here reflects insufficient task difficulty, as performance at the most adverse SNR approached approximately 20%. Jamadar (2020) called for a reassessment of the CRUNCH model, given the limited number of studies that have directly tested or replicated its predictions. Altogether, these findings suggest that the demand-response function is broadly preserved with aging, with age-related differences reflected primarily in the magnitude rather than the shape of the response.

Instead, we propose that, rather than reflecting the progressive mobilization of cognitive resources, the inverted U-shaped profile may reflect a transition from increasing task engagement at moderate levels of difficulty to reduced engagement as listening demands become more extreme. Both the FUEL framework (Pichora-Fuller et al., 2016) and the MoLE model (Herrmann and Johnsrude, 2020) draw an important distinction between resource recruitment and engagement. In these models, task engagement is defined as the active and motivated orientation of the listening process toward a valued communicative goal, the maintenance of which depends not only on the demands of the task, but also on the listener’s ongoing assessment of the relevance of continuing their efforts. Importantly, motivation and resource mobilization may be dissociable: a listener may be willing to invest effort regardless of whether the necessary cognitive resources are effectively recruited. Thus, DLPFC activity may track the degree to which listeners attempt to engage with the task rather than the degree to which they succeed. Supporting this view, Paul et al. (2021) showed that the quadratic component of parietal alpha power during speech-in-noise was specifically accounted for by subjective effort ratings rather than objective performance, and was preserved when analysis was restricted to correctly answered trials. These results suggest that the non-linear neural patterns during listening may reflect the listener’s subjective assessment of perceived difficulty rather than behavioural success. Consistent with this, the quadratic modulation of DLPFC activity observed here was similarly preserved regardless of whether trials were answered correctly or incorrectly. Our results further showed that at the most adverse SNRs, HbO responses declined alongside a marked drop in behavioural accuracy, and the brain-behaviour relationship present across easier conditions was no longer significant. Once successful comprehension becomes sufficiently unlikely, the neural cost of sustained engagement may diminish, and prefrontal activity may no longer predict behavioural outcomes.

If this descending portion truly reflects motivational withdrawal, one might have expected older adults to disengage earlier given their greater listening difficulties. However, as seen in Figure 2, the rate of accuracy decline across SNR conditions was comparable across age groups. Similarly to the pattern of brain activity, differences were observed in the magnitude of overall performance rather than in the shape of the function. This may indicate that withdrawal occurred at similar points along the difficulty continuum regardless of age. However, direct measures of engagement and motivation would be needed to confirm this interpretation.

Although this engagement-based interpretation is theoretically consistent with our findings, it should be viewed cautiously. The quadratic effect was modest in magnitude and may partly reflect large inter-individual variability in engagement. This small effect size may also be partly due to the design of the task. The mixed design used here reduced the predictability of trial difficulty, which may have encouraged participants to remain engaged even under highly challenging conditions, thereby reducing the withdrawal effect. A blocked design incorporating a clearly unachievable condition would more plausibly elicit stronger, more consistent disengagement responses. Furthermore, future studies should incorporate self-report measures that can explicitly capture task withdrawal, since the listening effort scale employed in this study cannot distinguish between continued engagement that ultimately fails and the decision to disengage altogether. Adapting the scale to include a withdrawal option or assessing motivation separately following blocks of differing difficulty would help to clarify this distinction.

### 4.2. DLPFC Activation and Neural Inefficiency in Older Adults

Our second hypothesis predicted that the functional significance of DLPFC recruitment would vary as a function of listening demand, such that increased activation would initially reflect compensatory recruitment before transitioning toward neural inefficiency and task disengagement. Contrary to this prediction, no evidence for compensatory recruitment was observed at any level of listening demand. Instead, greater DLPFC activation was consistently associated with poorer speech-in-noise performance across the ascending and peak portions of the demand-response function. These results are more consistent with the interpretation that increased DLPFC recruitment reflects neural inefficiency rather than successful compensation during effortful listening. Importantly, this negative brain-behaviour relationship was no longer evident under the most adverse listening conditions, consistent with the engagement-based interpretation proposed above and suggesting a transition from neural inefficiency to task disengagement as listening demands became extreme.

These findings extend our previous work (Perron et al., 2025), in which greater DLPFC activation was likewise associated with poorer behavioural performance in an independent sample. They also align with a growing body of evidence suggesting that increased prefrontal recruitment in aging does not support successful speech-in-noise perception. Bak et al. (2025), for example, observed a comparable pattern during a dual-task listening scenario involving a driving simulation in older adults with hearing loss. Likewise, hearing aid amplification has been shown to improve speech-in-noise performance while simultaneously reducing prefrontal activation (Vaisberg et al., 2025; Vaisberg et al., 2024), further suggesting that elevated DLPFC recruitment reflects inefficient processing under degraded listening conditions rather than beneficial adaptation.

Additional support for neural inefficiency comes from the subgroup analyses. High-performing older adults showed DLPFC activation profiles that closely resembled those of younger adults while also demonstrating comparable behavioural accuracy and listening effort across SNR conditions. In contrast, low-performing older adults showed substantially greater bilateral DLPFC recruitment together with poorer behavioural performance. The convergence of neural activity and behavioural performance, together with the similarity between younger adults and high-performing older adults across the neural, behavioural, and subjective measures, suggests that successful speech-in-noise perception in aging depends less on recruiting additional frontal resources than on preserving a more efficient, youth-like pattern of neural processing. This interpretation is consistent with the brain maintenance framework (Cabeza et al., 2018), which proposes that successful cognitive aging is characterized by the preservation of youthful neural organization and functional dynamics. Nevertheless, these subgroup findings should be interpreted cautiously because dividing the older adult sample further reduced statistical power.

The observed individual differences in neural inefficiency may reflect underlying differences in neural integrity. Importantly, we do not interpret elevated frontal recruitment as an effortful strategy that could simply be reduced to improve performance. Rather, we view it as a potential downstream consequence of long-standing degradation of auditory input: reduced fidelity of the afferent signal may place greater and more sustained demands on central mechanisms responsible for extracting task-relevant information from noise, contributing over time to less efficient neural processing. Several age-related neural changes could contribute to this process. One possibility is that age-related reductions in inhibitory neural circuitry reduce the ability to suppress task-irrelevant activity (Lalwani et al., 2019). Under this account, cortical regions that are effectively downregulated in younger adults become increasingly disinhibited with age, producing greater but functionally inefficient activation. Consistent with this possibility, we observed that younger adults showed relative DLPFC downregulation whereas older adults exhibited clear DLPFC upregulation. A second possibility is that age-related reductions in white matter integrity compromise the efficiency of large-scale neural communication, leading to less coordinated recruitment of prefrontal resources (Bennett and Rypma, 2013). Although white matter integrity was not measured in the present study, previous work has linked age-related degradation of prefrontal white matter pathways to poorer speech-in-noise perception (Perron et al., 2021; Tremblay et al., 2019). The present data cannot distinguish between these mechanisms, but both provide plausible explanations for the individual differences in DLPFC recruitment observed across older adults.

Consistent with these proposed mechanisms, chronological age was the strongest predictor of increased DLPFC activation within the older adult group. This finding is consistent with the possibility that neural inefficiency reflects progressive changes in brain structure and function, such that greater chronological age is associated with more pronounced DLPFC upregulation. This relationship emerged only at intermediate levels of difficulty, when the task was neither too easy nor too difficult, suggesting that these age-related neural changes become most apparent under substantial, but not overwhelming, listening demands. In contrast, hearing thresholds and global cognitive functioning were not significant predictors after correction for multiple comparisons, and including MoCA and better ear PTA as covariates did not alter the selected models. Together, these findings suggest that increased DLPFC recruitment is more closely linked to central neural aging than to peripheral hearing loss or global cognitive status, consistent with the evidence presented in the introduction that speech-in-noise difficulties extend beyond these factors.

Importantly, however, chronological age did not explain the preserved neural and behavioural profiles observed in high-performing older adults, as the two older adult subgroups did not significantly differ in age. Instead, these findings suggest that additional reserve-related factors likely determine whether individuals maintain a more youth-like pattern of neural recruitment. In line with the Scaffolding Theory of Aging and Cognition (Park and Reuter-Lorenz, 2009), factors such as lifelong cognitive engagement, physical activity, and social participation have been proposed to support the preservation of neural function with aging. Zhang et al. (2025) recently reported that older musicians exhibited both enhanced speech-in-noise perception and more youth-like frontotemporal connectivity. Although such reserve-related factors were not assessed in the present study, they represent plausible contributors.

A final aspect of the present findings concerns hemispheric lateralization. Although DLPFC recruitment in older adults was broadly bilateral, we observed that its relationship with performance was more pronounced in the left hemisphere, with greater left-lateralized activity observed in lower-performing older adults. This pattern is consistent with prior work implicating left prefrontal regions in effortful speech processing (for meta-analyses, see Alain et al., 2018; Perron et al., 2024). At the same time, it differs from our previous findings, in which performance-related effects were more right-lateralized (Perron et al., 2025). The reasons for this difference remain unclear. It may reflect differences in task design. Differences in participant characteristics may also have contributed, as the current sample included individuals with greater degrees of hearing loss than in our previous study. Vaisberg et al. reported that hearing aid amplification in older adults with hearing loss was associated with reduced DLPFC activity, with effects observed either bilaterally (Vaisberg et al., 2024) or predominantly in the left hemisphere (Vaisberg et al., 2024). Nevertheless, across studies, a consistent observation is that DLPFC activity tracks listening difficulty, supporting its role as a marker of effortful processing even if its precise hemispheric expression varies across paradigms and requires further investigation.

### 4.3. The Role of VLPFC and DMPFC

We conducted exploratory analyses on the VLPFC and DMPFC. We found a significant modulation of HbO depending on SNR or age in both regions, but HbO responses were unrelated to accuracy in either region. Therefore, our interpretations below focus on task-modulated engagement rather than direct behavioural relevance.

HbO responses in the VLPFC followed a strong quadratic relationship with SNR that closely resembled the pattern observed in the DLPFC. However, unlike the DLPFC, VLPFC responses were not modulated by age or any other demographic or behavioural variable. Importantly, the fNIRS montage used in the present study mostly captured activity from the posterior inferior frontal gyrus. Within a predictive coding framework, this profile is consistent with a role for the posterior inferior frontal gyrus in generating and maintaining high-level contextual priors, including phonological, lexical, and semantic expectations, that constrain perceptual inference across distributed auditory networks (Cope et al., 2017; Davis and Johnsrude, 2007). The quadratic effect may therefore reflect maximal engagement of predictive mechanisms at intermediate levels of speech degradation, where the acoustic signal remains sufficiently informative to support contextual inference but insufficiently clear to permit effortless comprehension. At the extremes of the SNR continuum, predictive support may be less necessary during clear speech and less effective when speech becomes severely degraded. The absence of age-related modulation is also consistent with evidence suggesting predictive coding mechanisms supporting speech comprehension may be relatively preserved in aging. Rysop et al. (2022) found that when stimulus intelligibility was calibrated to individual hearing abilities, older and younger adults recruited a comparable set of left-hemispheric language regions, and showed an equivalent behavioural predictability gain. In that study, age-related differences did not manifest at the level of regional activation but rather in the effective connectivity among recruited regions.

HbO responses in the DMPFC showed a distinct profile characterized by a significant main effect of age and a negative relationship between activity and SNR. This pattern is consistent with the established role of the DMPFC as a core hub of the default mode network, which is typically downregulated during externally oriented, demanding tasks (McKiernan et al., 2003). The progressive suppression of DMPFC activity with decreasing SNR likely reflects sustained allocation of attentional resources toward the auditory task and a corresponding disengagement from internally oriented processes. At first glance, the continued suppression of DMPFC activity under increasingly adverse SNRs may appear inconsistent with the interpretation that the descending portion of the DLPFC response reflects motivational withdrawal. However, these effects may index partially distinct components of task engagement. DMPFC suppression may reflect sustained externally directed attention and reduced internally oriented processing, whereas the downturn in DLPFC activity may reflect a reduction in the active recruitment of executive control resources once additional effort is no longer perceived as beneficial. Under this interpretation, listeners may remain externally oriented toward the task while simultaneously reducing costly top-down control processes.

The observed age effect in the DMPFC aligns with prior evidence indicating reduced efficiency in suppressing default mode activity in older adults during challenging cognitive tasks (Grady et al., 2016; Sambataro et al., 2010). Along these lines, the present findings suggest that age-related differences in speech-in-noise perception may also reflect a reduced capacity to suppress internally oriented processes as task demands increase. As Grady et al. (2006) proposed, alterations in the balance between default-mode and task-related activity may heighten susceptibility to distraction from irrelevant information during cognitive tasks.

### 4.4. Limitations and Future Directions

Several limitations and areas for future research must be acknowledged. First, although the overall sample size was comparable to previous fNIRS studies, the brain-behaviour analyses were conducted separately within each age group, resulting in relatively modest sample sizes. Consequently, these associations may be sensitive to sampling variability and should be interpreted cautiously. Replication in larger cohorts will be important to establish the stability of the observed relationships and to determine more confidently whether they distinguish between competing theoretical accounts of age-related prefrontal recruitment. Larger cohorts will also facilitate investigation of additional factors that were not assessed here, such as dietary habits, cholesterol levels, cognitive reserve, musical expertise, physical activity and social engagement, all of which may meaningfully contribute to variability in neural and behavioural outcomes. This could help to explain why some older adults maintain high performance and identify lifestyles and cognitive profiles that confer resilience against age-related neural inefficiency.

Secondly, although the present study expanded the coverage of fNIRS to include a larger portion of the prefrontal cortex than in our previous work, the spatial specificity of the observed effects remains limited by the spatial resolution of fNIRS. It is not possible to determine the precise anatomical localisation of the inefficiency pattern with the current montage, but it is likely to be localised within the middle frontal gyrus (see Supplementary Material 2). To overcome this challenge, approaches with a higher spatial resolution would be necessary, such as functional fMRI or higher density fNIRS configurations. Nevertheless, our results suggest that the signal of interest can be captured using a relatively accessible and portable method, which has important implications for measurements in applied and clinical contexts.

Thirdly, although stimulus presentation levels were individualized to reduce variability arising from differences in basic audibility, this procedure does not equate all aspects of auditory function across participants. Age-related suprathreshold deficits, such as reduced temporal processing, poorer frequency selectivity, or altered loudness growth, may still have contributed to speech-in-noise perception and associated neural responses.

Finally, and perhaps most importantly, our design was not optimised to investigate the underlying biological mechanisms. Multimodal imaging approaches that combine fNIRS or fMRI with measures of neurotransmitter function, white matter integrity, grey matter morphology, and inhibitory dynamics would help disentangle the neural substrates and advance our mechanistic understanding of how the aging brain adapts to increasing listening demands.

### 4.5. Conclusions

The present findings suggest that two partially dissociable properties of the DLPFC response contribute to speech-in-noise perception in aging. The overall shape of the demand-response trajectory may reflect changes in task engagement across increasing listening difficulty, whereas the magnitude of activation along that trajectory may index neural processing efficiency and predict behavioural success. This study provides further evidence that age-related activation of the DLPFC is more consistent with a pattern of neural inefficiency than with a pattern of compensation. Additionally, this effect was not explained by hearing thresholds or global cognitive status. Finally, we observed that high-performing older adults showed a demand-sensitive profile of behavioural performance and DLPFC recruitment that more closely resembling that of younger adults, supporting that preserved neural efficiency may distinguish resilient from vulnerable aging. From a clinical perspective, the ability to capture these neural signatures using fNIRS, an inexpensive and accessible technology, opens promising avenues for the early identification of at-risk individuals and the development of targeted interventions. Future research should focus on clarifying the mechanisms underlying age-related DLFC upregulation activity and identifying the factors that support preserved listening-related brain function in aging.

## Acknowledgements

We thank all participants for their time and commitment. This work was supported by a Discovery Grant from the Natural Sciences and Engineering Research Council of Canada (NSERC), awarded to F.A.R. (RGPIN-2023-03897). M.P. received a Postdoctoral Fellowship from the Canadian Institutes of Health Research.

## 5. CReDiT

**M.P.** Conceptualization, Data curation, Formal analysis, Investigation, Methodology, Visualization, Writing – original draft**. F.A.R.** Conceptualization, Funding acquisition, Project administration, Resources, Supervision, Validation, Writing – review and edition.

## 7. Data Statement

Raw behavioural and fNIRS data are available in the Borealis database (https://doi.org/10.5683/SP3/JSMAYV).

## References

Alain, C., Du, Y., Bernstein, L.J., Barten, T., Banai, K., 2018. Listening under difficult conditions: An activation likelihood estimation meta-analysis. Hum Brain Mapp 39(7), 2695–2709. 10.1002/hbm.24031.

Bak, K., Darakjian, L., Russo, F.A., Pichora-Fuller, M.K., Campos, J.L., 2025. Dual-task costs of listening while driving in older adults with and without audiometric hearing loss: Behavioural and neurophysiological outcomes. Hear Res 468, 109437. 10.1016/j.heares.2025.109437.

Bennett, I.J., Rypma, B., 2013. Advances in functional neuroanatomy: a review of combined DTI and fMRI studies in healthy younger and older adults. Neurosci Biobehav Rev 37(7), 1201–1210. 10.1016/j.neubiorev.2013.04.008.

Bilodeau-Mercure, M., Lortie, C.L., Sato, M., Guitton, M.J., Tremblay, P., 2015. The neurobiology of speech perception decline in aging. Brain Struct Funct 220(2), 979–997. 10.1007/s00429-013-0695-3.

Bramhall, N.F., McMillan, G.P., 2024. Perceptual Consequences of Cochlear Deafferentation in Humans. Trends Hear 28, 23312165241239541. 10.1177/23312165241239541.

Burnham, K.P., Anderson, D.R., 2002. Model selection and multimodel inference: a practical information-theoretic approach. Springer.

Cabeza, R., Albert, M., Belleville, S., Craik, F.I.M., Duarte, A., Grady, C.L., Lindenberger, U., Nyberg, L., Park, D.C., Reuter-Lorenz, P.A., Rugg, M.D., Steffener, J., Rajah, M.N., 2018. Maintenance, reserve and compensation: the cognitive neuroscience of healthy ageing. Nat Rev Neurosci 19(11), 701–710. 10.1038/s41583-018-0068-2.

Cope, T.E., Sohoglu, E., Sedley, W., Patterson, K., Jones, P.S., Wiggins, J., Dawson, C., Grube, M., Carlyon, R.P., Griffiths, T.D., Davis, M.H., Rowe, J.B., 2017. Evidence for causal top-down frontal contributions to predictive processes in speech perception. Nat Commun 8(1), 2154. 10.1038/s41467-017-01958-7.

Cui, X., Bray, S., Bryant, D.M., Glover, G.H., Reiss, A.L., 2011. A quantitative comparison of NIRS and fMRI across multiple cognitive tasks. Neuroimage 54(4), 2808–2821. 10.1016/j.neuroimage.2010.10.069.

Davis, M.H., Johnsrude, I.S., 2007. Hearing speech sounds: top-down influences on the interface between audition and speech perception. Hear Res 229(1-2), 132–147. 10.1016/j.heares.2007.01.014.

Du, Y., Buchsbaum, B.R., Grady, C.L., Alain, C., 2016. Increased activity in frontal motor cortex compensates impaired speech perception in older adults. Nat Commun 7, 12241. 10.1038/ncomms12241.

Fabrizio-Stover, E.M., Dias, J.W., McClaskey, C.M., Harris, K.C., 2026. Age-related auditory nerve deficits propagate central gain throughout the auditory system: Associations with cortical microstructure and speech recognition. Neurobiol Aging 157, 98–110. 10.1016/j.neurobiolaging.2025.10.007.

Fostick, L., Ben-Artzi, E., Babkoff, H., 2013. Aging and speech perception: beyond hearing threshold and cognitive ability. J Basic Clin Physiol Pharmacol 24(3), 175–183. 10.1515/jbcpp-2013-0048.

Fullgrabe, C., Moore, B.C., Stone, M.A., 2014. Age-group differences in speech identification despite matched audiometrically normal hearing: contributions from auditory temporal processing and cognition. Front Aging Neurosci 6, 347. 10.3389/fnagi.2014.00347.

Grady, C., Sarraf, S., Saverino, C., Campbell, K., 2016. Age differences in the functional interactions among the default, frontoparietal control, and dorsal attention networks. Neurobiol Aging 41, 159–172. 10.1016/j.neurobiolaging.2016.02.020.

Grady, C.L., Springer, M.V., Hongwanishkul, D., McIntosh, A.R., Winocur, G., 2006. Age-related changes in brain activity across the adult lifespan. J Cogn Neurosci 18(2), 227–241. 10.1162/089892906775783705.

Herrmann, B., Johnsrude, I.S., 2020. A model of listening engagement (MoLE). Hear Res 397, 108016. 10.1016/j.heares.2020.108016.

Humes, L.E., 2019. The World Health Organization’s hearing-impairment grading system: an evaluation for unaided communication in age-related hearing loss. Int J Audiol 58(1), 12–20. 10.1080/14992027.2018.1518598.

Huppert, T.J., Diamond, S.G., Franceschini, M.A., Boas, D.A., 2009. HomER: a review of time-series analysis methods for near-infrared spectroscopy of the brain. Appl. Opt. 48(10), D280–D298. 10.1364/AO.48.00D280.

Jamadar, S.D., 2020. The CRUNCH model does not account for load-dependent changes in visuospatial working memory in older adults. Neuropsychologia 142, 107446. 10.1016/j.neuropsychologia.2020.107446.

Kwasa, J., Peterson, H.M., Karrobi, K., Jones, L., Parker, T., Nickerson, N., Wood, S., 2023. Demographic reporting and phenotypic exclusion in fNIRS. Front Neurosci 17, 1086208. 10.3389/fnins.2023.1086208.

Lalwani, P., Gagnon, H., Cassady, K., Simmonite, M., Peltier, S., Seidler, R.D., Taylor, S.F., Weissman, D.H., Polk, T.A., 2019. Neural distinctiveness declines with age in auditory cortex and is associated with auditory GABA levels. Neuroimage 201, 116033. 10.1016/j.neuroimage.2019.116033.

Lawrence, R.J., Wiggins, I.M., Anderson, C.A., Davies-Thompson, J., Hartley, D.E.H., 2018. Cortical correlates of speech intelligibility measured using functional near-infrared spectroscopy (fNIRS). Hear Res 370, 53–64. 10.1016/j.heares.2018.09.005.

Lin, V.Y., Chung, J., Callahan, B.L., Smith, L., Gritters, N., Chen, J.M., Black, S.E., Masellis, M., 2017. Development of cognitive screening test for the severely hearing impaired: Hearing-impaired MoCA. Laryngoscope 127 Suppl 1, S4–s11. 10.1002/lary.26590.

Logan, J.M., Sanders, A.L., Snyder, A.Z., Morris, J.C., Buckner, R.L., 2002. Under-recruitment and nonselective recruitment: dissociable neural mechanisms associated with aging. Neuron 33(5), 827–840. 10.1016/s0896-6273(02)00612-8.

McKiernan, K.A., Kaufman, J.N., Kucera-Thompson, J., Binder, J.R., 2003. A parametric manipulation of factors affecting task-induced deactivation in functional neuroimaging. J Cogn Neurosci 15(3), 394–408. 10.1162/089892903321593117.

Park, D.C., Reuter-Lorenz, P., 2009. The adaptive brain: aging and neurocognitive scaffolding. Annu Rev Psychol 60, 173–196. 10.1146/annurev.psych.59.103006.093656.

Paul, B.T., Chen, J., Le, T., Lin, V., Dimitrijevic, A., 2021. Cortical alpha oscillations in cochlear implant users reflect subjective listening effort during speech-in-noise perception. PLoS One 16(7), e0254162. 10.1371/journal.pone.0254162.

Peelle, J.E., 2018. Listening Effort: How the Cognitive Consequences of Acoustic Challenge Are Reflected in Brain and Behavior. Ear Hear 39(2), 204–214. 10.1097/aud.0000000000000494.

Perron, M., Shatzer, H., Zara, M., Russo, F., 2025. Age-related increased frontal activation in sentence comprehension reflects inefficiency, not compensation. Neurobiol Aging 155, 100–112. 10.1016/j.neurobiolaging.2025.07.014.

Perron, M., Theaud, G., Descoteaux, M., Tremblay, P., 2021. The frontotemporal organization of the arcuate fasciculus and its relationship with speech perception in young and older amateur singers and non-singers. Hum Brain Mapp n/a(n/a), 1–19. 10.1002/hbm.25416.

Perron, M., Vuong, V., Grassi, M.W., Imran, A., Alain, C., 2024. Engagement of the speech motor system in challenging speech perception: Activation likelihood estimation meta-analyses. Hum Brain Mapp 45(13), e70023. 10.1002/hbm.70023.

Pichora-Fuller, M.K., Kramer, S.E., Eckert, M.A., Edwards, B., Hornsby, B.W., Humes, L.E., Lemke, U., Lunner, T., Matthen, M., Mackersie, C.L., Naylor, G., Phillips, N.A., Richter, M., Rudner, M., Sommers, M.S., Tremblay, K.L., Wingfield, A., 2016. Hearing Impairment and Cognitive Energy: The Framework for Understanding Effortful Listening (FUEL). Ear Hear 37 Suppl 1, 5s–27s. 10.1097/aud.0000000000000312.

Plichta, M.M., Herrmann, M.J., Baehne, C.G., Ehlis, A.C., Richter, M.M., Pauli, P., Fallgatter, A.J., 2006. Event-related functional near-infrared spectroscopy (fNIRS): are the measurements reliable? Neuroimage 31(1), 116–124. 10.1016/j.neuroimage.2005.12.008.

Reuter-Lorenz, P.A., Cappell, K.A., 2008. Neurocognitive Aging and the Compensation Hypothesis. Curr. Dir. Psychol. Sci 17(3), 177–182. 10.1111/j.1467-8721.2008.00570.x.

Reuter-Lorenz, P.A., Marshuetz, C., Jonides, J., Smith, E.E., Hartley, A., Koeppe, R., 2001. Neurocognitive ageing of storage and executive processes. Eur. J. Cogn. Psychol 13(1-2), 257–278.

RStudio Team, 2022. RStudio: Integrated Development Environment for R. RStudio, PBC, Boston, MA.

Russo, F.A., Pichora-Fuller, M.K., 2008. Tune in or tune out: age-related differences in listening to speech in music. Ear Hear 29(5), 746–760. 10.1097/AUD.0b013e31817bdd1f.

Ryan, D.B., Eckert, M.A., Sellers, E.W., Schairer, K.S., McBee, M.T., Jones, M.R., Smith, S.L., 2022. Impact of Effortful Word Recognition on Supportive Neural Systems Measured by Alpha and Theta Power. Ear Hear 43(5), 1549–1562. 10.1097/aud.0000000000001211.

Rysop, A.U., Schmitt, L.-M., Obleser, J., Hartwigsen, G., 2022. Age-related differences in the neural network interactions underlying the predictability gain. Cortex 154, 269–286. 10.1016/j.cortex.2022.05.020.

Sambataro, F., Murty, V.P., Callicott, J.H., Tan, H.Y., Das, S., Weinberger, D.R., Mattay, V.S., 2010. Age-related alterations in default mode network: impact on working memory performance. Neurobiol Aging 31(5), 839–852. 10.1016/j.neurobiolaging.2008.05.022.

Stevens, G., Flaxman, S., Brunskill, E., Mascarenhas, M., Mathers, C.D., Finucane, M., 2013. Global and regional hearing impairment prevalence: an analysis of 42 studies in 29 countries. Eur J Public Health 23(1), 146–152. 10.1093/eurpub/ckr176.

Tadel, F., Baillet, S., Mosher, J.C., Pantazis, D., Leahy, R.M., 2011. Brainstorm: a user-friendly application for MEG/EEG analysis. Comput Intell Neurosci 2011, 879716. 10.1155/2011/879716.

Tremblay, P., Perron, M., Deschamps, I., Kennedy-Higgins, D., Houde, J.C., Dick, A.S., Descoteaux, M., 2019. The role of the arcuate and middle longitudinal fasciculi in speech perception in noise in adulthood. Hum Brain Mapp 40(1), 226–241. 10.1002/hbm.24367.

Vaisberg, J.M., Dang, C., Jiang, Y., Qian, J., Russo, F.A., 2025. Brain benefits of deep learning-based noise management in experienced hearing aid users using functional near infrared spectroscopy. Sci Rep 15(1), 41815. 10.1038/s41598-025-25801-y.

Vaisberg, J.M., Gilmore, S., Qian, J., Russo, F.A., 2024. The Benefit of Hearing Aids as Measured by Listening Accuracy, Subjective Listening Effort, and Functional Near Infrared Spectroscopy. Trends Hear 28, 23312165241273346. 10.1177/23312165241273346.

Wong, P.C., Jin, J.X., Gunasekera, G.M., Abel, R., Lee, E.R., Dhar, S., 2009. Aging and cortical mechanisms of speech perception in noise. Neuropsychologia 47(3), 693–703. 10.1016/j.neuropsychologia.2008.11.032.

Wu, Y.H., Stangl, E., Zhang, X., Perkins, J., Eilers, E., 2016. Psychometric Functions of Dual-Task Paradigms for Measuring Listening Effort. Ear Hear 37(6), 660–670. 10.1097/aud.0000000000000335.

Yücel, M.A., Lühmann, A.V., Scholkmann, F., Gervain, J., Dan, I., Ayaz, H., Boas, D., Cooper, R.J., Culver, J., Elwell, C.E., Eggebrecht, A., Franceschini, M.A., Grova, C., Homae, F., Lesage, F., Obrig, H., Tachtsidis, I., Tak, S., Tong, Y., Torricelli, A., Wabnitz, H., Wolf, M., 2021. Best practices for fNIRS publications. Neurophotonics 8(1), 012101. 10.1117/1.NPh.8.1.012101.

Zekveld, A.A., Kramer, S.E., 2014. Cognitive processing load across a wide range of listening conditions: insights from pupillometry. Psychophysiology 51(3), 277–284. 10.1111/psyp.12151.

Zhang, L., Ross, B., Du, Y., Alain, C., 2025. Long-term musical training can protect against age-related upregulation of neural activity in speech-in-noise perception. PLoS Biol 23(7), e3003247. 10.1371/journal.pbio.3003247.

Zimeo Morais, G.A., Balardin, J.B., Sato, J.R., 2018. fNIRS Optodes’ Location Decider (fOLD): a toolbox for probe arrangement guided by brain regions-of-interest. Sci Rep 8(1), 3341. 10.1038/s41598-018-21716-z.

