## Supplementary Material for "Non-Linear Dorsolateral Prefrontal Recruitment During Speech-in-Noise Perception in Aging"

### Su**pplementary Material 1.** Participant Characteristics Relevant to fNIRS Signal Acquisition.

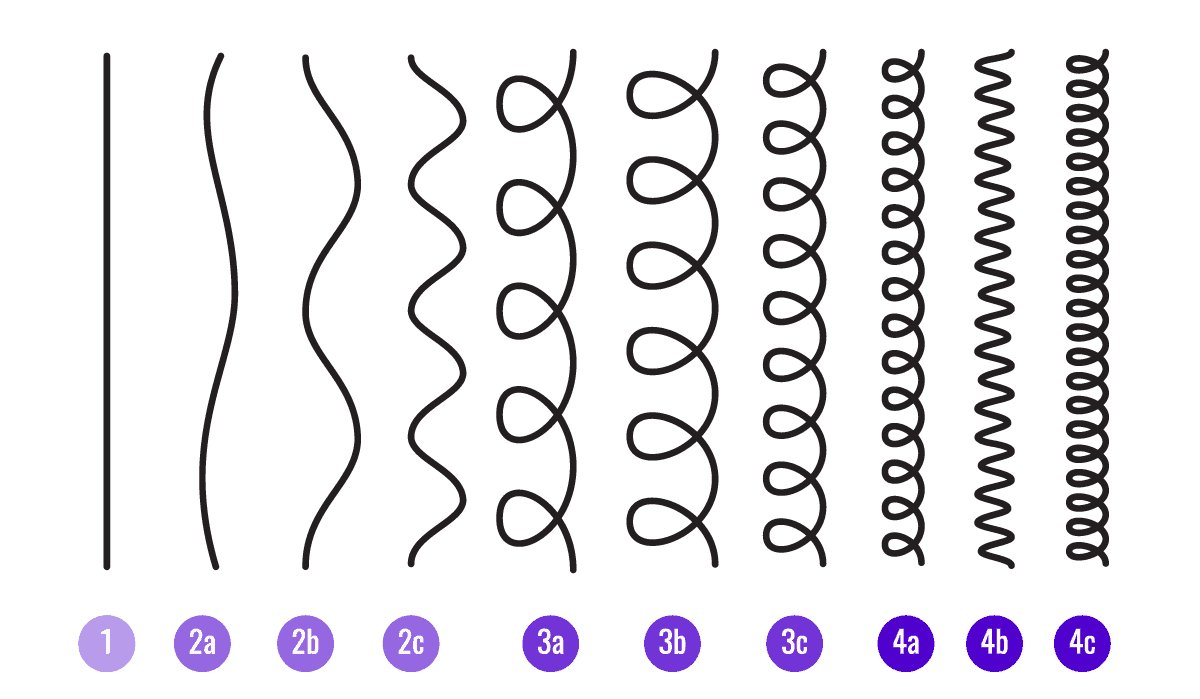

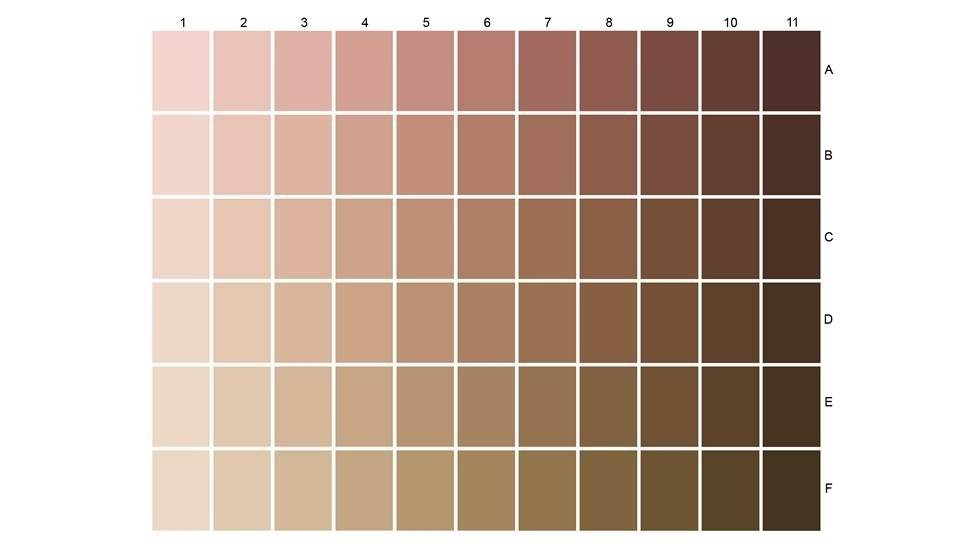

**L'Oreal Skin Color Chart	Andre Walker Hair Typing System**

Distribution of skin tones. Values not in brackets represent young adults, and values in brackets represent older adults.

| **1** | **2** | **3** | **4** | **5** | **6** | **7** | **8** | **9** | **10** |  |
| --- | --- | --- | --- | --- | --- | --- | --- | --- | --- | --- |
| 1 (2) | 1 (0) | 0 (1) | 0 (0) | 0 (0) | 1 (0) | 0 (0) | 0 (0) | 0 (0) | 0 (0) | **A** |
| 0 (1) | 3 (1) | 0 (1) | 0 (0) | 0 (0) | 0 (0) | 0 (0) | 0 (0) | 0 (0) | 0 (0) | **B** |
| 2 (5) | 3 (4) | 1 (3) | 0 (2) | 0 (0) | 3 (0) | 0 (0) | 0 (0) | 0 (1) | 0 (0) | **C** |
| 0 (3) | 1 (1) | 5 (1) | 0 (2) | 1 (0) | 1 (0) | 1 (1) | 0 (0) | 0 (0) | 1 (0) | **D** |
| 1 (0) | 3 (1) | 3 (2) | 1 (0) | 0 (1) | 0 (0) | 1 (0) | 0 (0) | 0 (0) | 0 (0) | **E** |
| 0 (0) | 0 (0) | 2 (0) | 0 (1) | 0 (0) | 0 (0) | 0 (0) | 0 (0) | 0 (0) | 0 (0) | **F** |

Distribution of hair types. Values not in brackets represent young adults, and values in brackets represent older adults.

| **1** | **2a** | **2b** | **2c** | **3a** | **3b** | **3c** | **4a** | **4b** | **4c** |
| --- | --- | --- | --- | --- | --- | --- | --- | --- | --- |
| 7 (11) | 9 (7) | 9 (7) | 6 (3) | 2 (3) | 1 () | 2 (2) | 0 () | 0 () | 0 (1) |

Head measurements.

| **Measure** | **Young Adults** | | **Older Adults** | |
| --- | --- | --- | --- | --- |
|  | **Mean** | **Standard Deviation** | **Mean** | **Standard Deviation** |
| Inion-nasion distance (cm) | 37.78 | 1.73 | 38.26 | 2.15 |
| Left-to-right periauricular area (cm) | 38.15 | 4.71 | 37.12 | 1.68 |
| Circumference (cm) | 60.28 | 3.87 | 60.53 | 1.57 |

Hair color. The values in brackets represent the percentage of the sample.

| **Hair color** | **Young adults** | **Older adults** |
| --- | --- | --- |
| Black | 14 (38.9%) | 3 (8.8%) |
| Brown (light and dark) | 19 (52.8%) | 5 (14.7%) |
| Blonde | 2 (5.6%) | 0 (0.0%) |
| Red | 1 (2.8%) | 2 (5.9%) |
| Grey/White* | 0 (0.0%) | 24 (70.6%) |

*Includes all hair described as grey, white, silver, or combinations of these with another color (e.g., grey/brown, grey/white, white/brown).

### **Supplementary Material 2.** Group-Averaged MNI Coordinates of Optode and Measurement Channels.

**Optode*.*** *Tx = transmitter; Rx = receiver.*

| **Optode** | **x** | **y** | **z** |
| --- | --- | --- | --- |
| Tx1 | 52 | 0 | 13 |
| Rx1 | 47 | 8 | -7 |
| Tx2 | 49 | 4 | -13 |
| Tx3 | 46 | 21 | 7 |
| Rx2 | 48 | 15 | 28 |
| Tx4 | 39 | 33 | 33 |
| Rx3 | 19 | 44 | 34 |
| Tx5 | 21 | 54 | 11 |
| Rx4 | 41 | 40 | 11 |
| Tx6 | -50 | 1 | 12 |
| Rx5 | -46 | 11 | -8 |
| Tx7 | -47 | 4 | -13 |
| Tx8 | -44 | 22 | 5 |
| Rx6 | -45 | 16 | 27 |
| Tx9 | -36 | 34 | 32 |
| Rx7 | -37 | 43 | 10 |
| Tx10 | -18 | 55 | 10 |
| Rx8 | -15 | 42 | 34 |

**Measurement channels.** *Tx = transmitter; Rx = receiver, SSC = Short separation channel.*

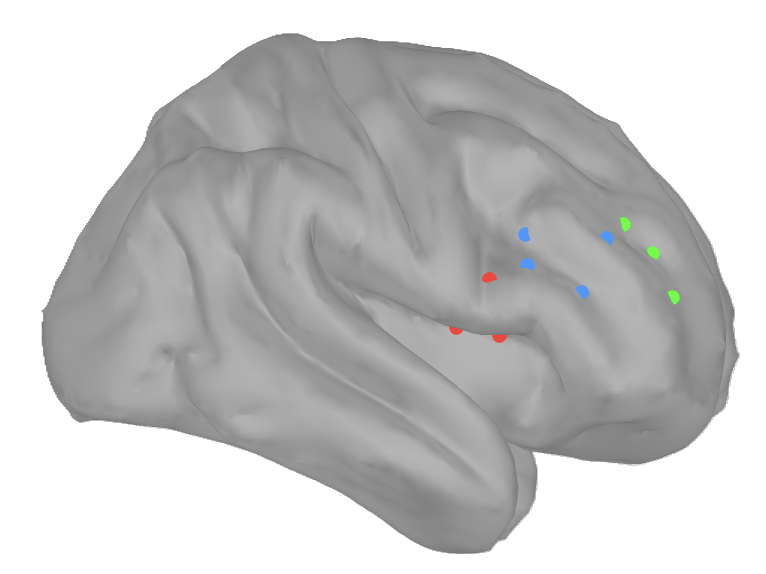

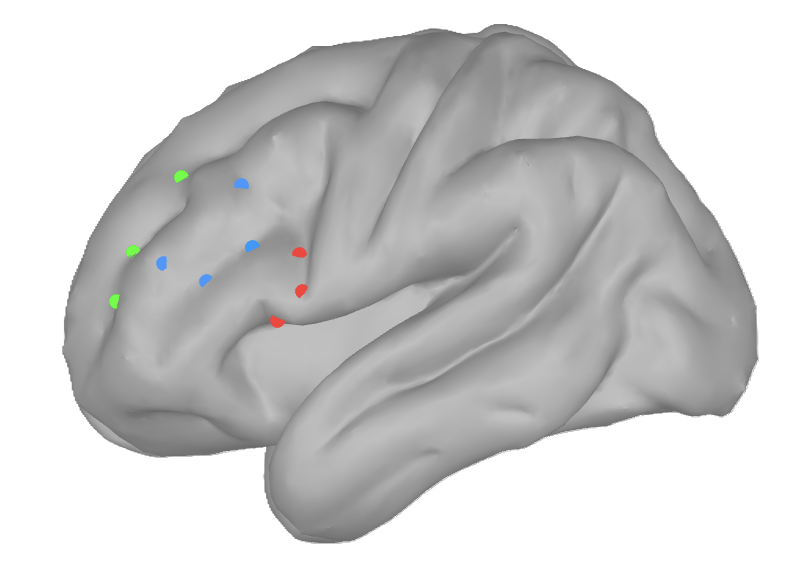

| **Hemisphere** | **Cluster** | **Channel** | **x** | **y** | **z** |
| --- | --- | --- | --- | --- | --- |
| Left | 1 | Tx6-Rx5 | -48 | 6 | 2 |
|  |  | Tx6-Rx6 | -48 | 9 | 19 |
|  |  | Tx8-Rx5 | -45 | 16 | -1 |
|  | 2 | Tx8-Rx6 | -45 | 19 | 16 |
|  |  | Tx8-Rx7 | -41 | 32 | 10 |
|  |  | Tx9-Rx6 | -40 | 25 | 29 |
|  |  | Tx9-Rx7 | -36 | 38 | 21 |
|  | 3 | Tx9-Rx8 | -25 | 38 | 33 |
|  |  | Tx10-Rx7 | -27 | 49 | 10 |
|  |  | Tx10-Rx8 | -17 | 49 | 22 |
|  | SSC | Tx7-Rx5 | -47 | 8 | -10 |
| Right | 1 | Tx1-Rx1 | 50 | 4 | 4 |
|  |  | Tx1-Rx2 | 50 | 8 | 20 |
|  |  | Tx3-Rx1 | 46 | 14 | 0 |
|  | 2 | Tx3-Rx2 | 47 | 18 | 17 |
|  |  | Tx3-Rx4 | 43 | 30 | 9 |
|  |  | Tx4-Rx2 | 43 | 24 | 30 |
|  |  | Tx4-Rx4 | 40 | 36 | 22 |
|  | 3 | Tx4-Rx3 | 29 | 38 | 34 |
|  |  | Tx5-Rx3 | 20 | 49 | 23 |
|  |  | Tx5-Rx4 | 31 | 47 | 11 |
|  | SSC | Tx2-Rx1 | 48 | 6 | -10 |

### **Supplementary Material 3.** Grand-Average Hemodynamic Response Functions (HRFs) Time-Locked to Stimulus Presentation for each Cluster, Signal-to-Noise Ratio Condition and Group.

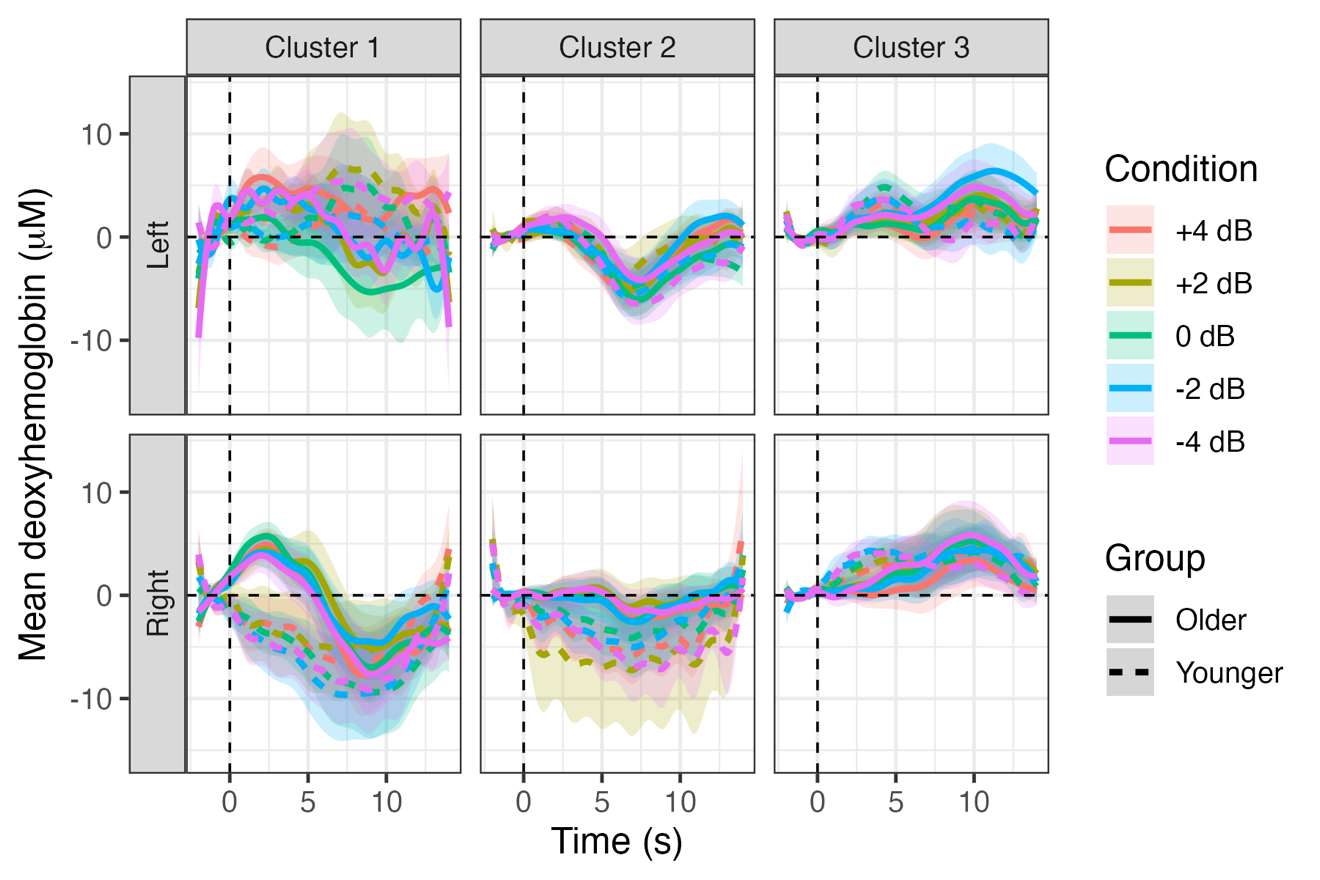

### **Supplementary 4.** Results of Deoxyhemoglobin (HbR) Analyses.

**HbR mixed-effects model results for the VLPFC *(cluster 1)* across OAs and YAs.**

The best-supported linear and hierarchical quadratic candidate sets converged on the same model, which included only Hemisphere as a fixed effect (HbR ~ Hemisphere + (1 | Participant)). Neither the linear nor quadratic SNR terms improved model fit, indicating no evidence for an association between HbR responses and listening difficulty.

| **Parameter** | **β** | **SE** | **95% CI** | **t** | **p** |
| --- | --- | --- | --- | --- | --- |
| (Intercept) | 0.11 | 0.09 | [-0.06, 0.28] | 1.26 | 0.208 |
| Hemisphere [Right] | -0.22 | 0.06 | [-0.34, -0.10] | -3.69 | **<0.001** |

**HbR mixed-effects model results for VLPFC *(cluster 1)* across OAs_L, OAs_H, and YAs.**

The best-supported linear and hierarchical quadratic candidate sets converged on the same model, which included only Hemisphere as a fixed effect (HbR ~ Hemisphere + (1 | Participant)). Neither the linear nor quadratic SNR terms improved model fit, indicating no evidence for an association between HbR responses and listening difficulty.

| **Parameter** | **β** | **SE** | **95% CI** | **t** | **p** |
| --- | --- | --- | --- | --- | --- |
| (Intercept) | 0.11 | 0.09 | [-0.06, 0.28] | 1.26 | 0.208 |
| Hemisphere [Right] | -0.22 | 0.06 | [-0.34, -0.10] | -3.69 | **<0.001** |

**HbR mixed-effects model results for the DLPFC *(cluster 2)* across OAs and YAs.**

The best-supported linear and hierarchical quadratic candidate sets converged on the same model, which included only Hemisphere as a fixed effect (HbR ~ Hemisphere + (1 | Participant)). Neither the linear nor quadratic SNR terms improved model fit, indicating no evidence for an association between HbR responses and listening difficulty.

| **Parameter** | **β** | **SE** | **95% CI** | **t** | **p** |
| --- | --- | --- | --- | --- | --- |
| (Intercept) | -0.05 | 0.09 | [-0.23, 0.13] | -0.54 | 0.590 |
| Hemisphere [Right] | 0.12 | 0.06 | [0.01, 0.23] | 2.12 | **0.035** |

**HbR mixed-effects model results for DLPFC *(cluster 2)* across OAs_L, OAs_H, and YAs.**

The best-supported linear and hierarchical quadratic candidate sets converged on the same model, which included Hemisphere, Group, and the Hemisphere × Group interaction (HbR ~ Hemisphere + Group + Hemisphere × Group + (1 | Participant)). Neither the linear nor quadratic SNR terms improved model fit, indicating no evidence for an association between HbR responses and listening difficulty.

| **Parameter** | **β** | **SE** | **95% CI** | **t** | **p** |
| --- | --- | --- | --- | --- | --- |
| (Intercept) | -0.03 | 0.13 | [-0.28, 0.21] | -0.28 | 0.782 |
| Hemisphere [Right] | 0.05 | 0.08 | [-0.10, 0.20] | 0.66 | 0.510 |
| Group [OAs_L] | 0.27 | 0.22 | [-0.16, 0.71] | 1.24 | 0.216 |
| Group [OAs_H] | -0.33 | 0.22 | [-0.77, 0.10] | -1.51 | 0.131 |
| Hemisphere [Right] x Group [OAs_L] | -0.07 | 0.13 | [-0.34, 0.19] | -0.55 | 0.585 |
| Hemisphere [Right] x Group [OAs_H] | 0.34 | 0.13 | [0.07, 0.60] | 2.52 | **0.012** |

**HbR mixed-effects model results for the DMPFC *(cluster 3)* across OAs and YAs.**

The best-supported linear and hierarchical quadratic candidate sets converged on the same null model (HbR ~ (1 | Participant)). Neither the linear nor quadratic SNR terms, nor Hemisphere, Group, or their interactions, improved model fit.

| **Parameter** | **β** | **SE** | **95% CI** | **t** | **p** |
| --- | --- | --- | --- | --- | --- |
| (Intercept) | 0.01 | 0.09 | [-0.18, 0.19] | 0.09 | 0.932 |

**HbR mixed-effects model results for DMPFC *(cluster 3)* across OAs_L, OAs_H, and YAs.**

The best-supported linear and hierarchical quadratic candidate sets converged on the same null model (HbR ~ (1 | Participant)). Neither the linear nor quadratic SNR terms, nor Hemisphere, Group, or their interactions, improved model fit.

| **Parameter** | **β** | **SE** | **95% CI** | **t** | **p** |
| --- | --- | --- | --- | --- | --- |
| (Intercept) | 0.01 | 0.09 | [-0.18, 0.19] | 0.09 | 0.932 |

### **Supplementary Material 5.** Results for Oxyhemoglobin (HbO) for the Ventrolateral Prefrontal Cortex (VLPFC) and the Dorsomedial Prefrontal Cortex (DMPFC).

**HbO mixed-effects model results for the VLPFC *(cluster 1)* across OAs and YAs.**

The best-supported linear model included the linear SNR term, Group, and the linear SNR × Group interaction (HbO ~ linear SNR + Group + linear SNR × Group + (1 | Participant)), whereas the best-supported hierarchical quadratic model additionally retained the quadratic SNR term (HbO ~ linear SNR + quadratic SNR + Group + linear SNR × Group + (1 | Participant)). Comparison of the candidate models indicated that the hierarchical quadratic model provided a better fit to the data (AICc = 6154.2 vs. 6157.4; ΔAICc = 3.23; Akaike weights = 0.834 and 0.166, respectively).

| **Parameter** | **β** | **SE** | **95% CI** | **t** | **p** |
| --- | --- | --- | --- | --- | --- |
| (Intercept) | -0.10 | 0.13 | [-0.37, 0.16] | -0.76 | 0.450 |
| SNR [Linear] | -0.11 | 0.03 | [-0.18, -0.04] | -3.09 | **0.002** |
| SNR [Quadratic] | -0.06 | 0.02 | [-0.10, -0.01] | -2.30 | **0.022** |
| Group [OAs] | 0.21 | 0.19 | [-0.16, 0.58] | 1.10 | 0.270 |
| SNR [Linear] x Group [OAs] | 0.09 | 0.05 | [0.00, 0.19] | 1.88 | 0.061 |

**HbO mixed-effects model results for VLPFC *(cluster 1)* across OAs_L, OAs_H, and YAs.**

The best-supported linear model included the linear SNR term, Hemisphere, Group, and the Group × Hemisphere interaction (HbO ~ linear SNR + Hemisphere + Group + Group × Hemisphere + (1 | Participant)), whereas the best-supported hierarchical quadratic model additionally retained the quadratic SNR term (HbO ~ linear SNR + quadratic SNR + Hemisphere + Group + Group × Hemisphere + (1 | Participant)). Comparison of the candidate models indicated that the hierarchical quadratic model provided a better fit to the data (AICc = 6143.4 vs. 6146.8; ΔAICc = 3.41; Akaike weights = 0.846 and 0.154, respectively).

| **Parameter** | **β** | **SE** | **95% CI** | **t** | **p** |
| --- | --- | --- | --- | --- | --- |
| (Intercept) | -0.09 | 0.14 | [-0.36, 0.18] | -0.65 | 0.513 |
| SNR [Linear] | -0.06 | 0.02 | [-0.11, -0.01] | -2.52 | **0.012** |
| SNR [Quadratic] | -0.06 | 0.02 | [-0.10, -0.01] | -2.35 | **0.019** |
| Hemisphere [Right] | -0.02 | 0.07 | [-0.16, 0.11] | -0.31 | 0.753 |
| Group [OAs_L] | 0.07 | 0.24 | [-0.40, 0.54] | 0.27 | 0.784 |
| Group [OAs_H] | 0.24 | 0.24 | [-0.23, 0.71] | 0.99 | 0.324 |
| Hemisphere [Right] x Group [OAs_L] | 0.41 | 0.12 | [0.18, 0.64] | 3.46 | **0.001** |
| Hemisphere [Right] x Group [OAs_H] | -0.18 | 0.12 | [-0.41, 0.06] | -1.49 | 0.137 |

Group differences in HbO concentration. (A) Older adults versus younger adults. (B) Young adults, low performing older adults (OA_L), and high performing older adults (OA_H). Shaded areas around the bars represent the standard error. Each point represents an individual participant.

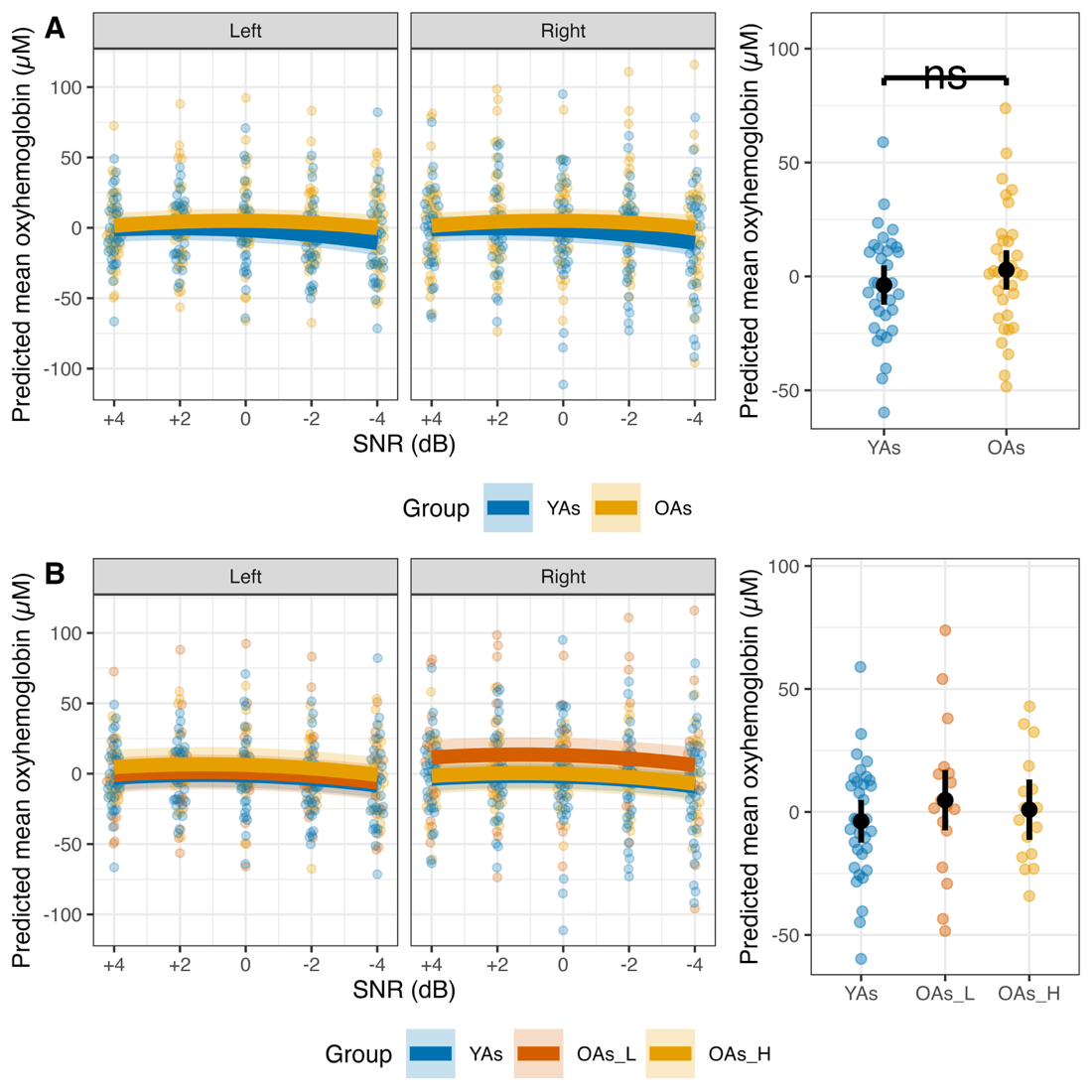

Correlation coefficients by age group and hemisphere. (A) Correlations between accuracy, self-reported listening effort, and HbO within each signal-to-noise ratio (SNR) and hemisphere. (B) Pearson correlations between HbO and demographic measures, including age, best-ear pure tone average (BestPTA), and MoCA scores. Asterisks indicate significant correlations.

*No significant correlations were found.*

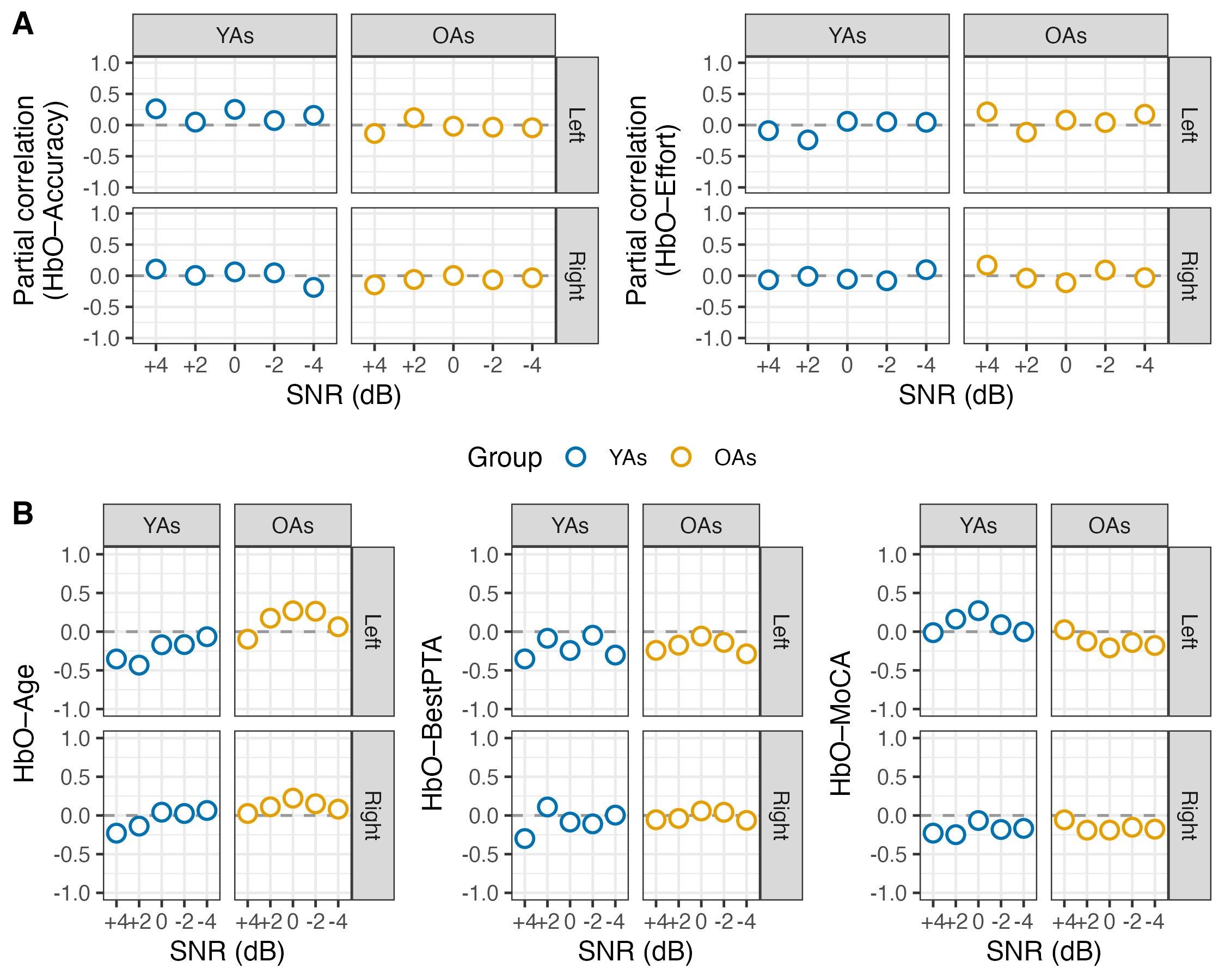

**HbO mixed-effects model results for the DMPFC *(cluster 3)* across OAs and YAs.**

The best-supported linear model included the linear SNR term and Group (HbO ~ linear SNR + Group + (1 | Participant)), whereas the best-supported hierarchical quadratic model additionally retained the quadratic SNR term (HbO ~ linear SNR + quadratic SNR + Group + (1 | Participant)). Comparison of the candidate models indicated that the linear model provided the better fit to the data (AICc = 6049.4 vs. 6051.2; ΔAICc = 1.81; Akaike weight = 0.712 vs. 0.288).

| **Parameter** | **β** | **SE** | **95% CI** | **t** | **p** |
| --- | --- | --- | --- | --- | --- |
| (Intercept) | -0.46 | 0.13 | [-0.73, -0.20] | -3.48 | **0.001** |
| SNR [Linear] | -0.05 | 0.02 | [-0.09, -0.01] | -2.54 | **0.011** |
| Group [OAs] | 0.90 | 0.19 | [0.53, 1.27] | 4.79 | **<0.001** |

**HbO mixed-effects model results for DMPFC *(cluster 3)* across OAs_L, OAs_H, and YAs.**

The best-supported linear model included the linear SNR term and Group (HbO ~ linear SNR + Group + (1 | Participant)), whereas the best-supported hierarchical quadratic model additionally retained the quadratic SNR term (HbO ~ linear SNR + quadratic SNR + Group + (1 | Participant)). Comparison of the candidate models indicated that the linear model provided the better fit to the data (AICc = 6048.9 vs. 6050.8; ΔAICc = 1.82; Akaike weight = 0.713 vs. 0.287).

| **Parameter** | **β** | **SE** | **95% CI** | **t** | **p** |
| --- | --- | --- | --- | --- | --- |
| (Intercept) | -0.46 | 0.13 | [-0.72, -0.21] | -3.54 | **<0.001** |
| SNR [Linear] | -0.05 | 0.02 | [-0.09, -0.01] | -2.54 | **0.011** |
| Group [OAs_L] | 1.11 | 0.23 | [0.67, 1.55] | 4.90 | **<0.001** |
| Group [OAs_H] | 0.69 | 0.23 | [0.25, 1.14] | 3.06 | **0.002** |

Group differences in HbO concentration. (A) Older adults versus younger adults. (B) Young adults, low performing older adults (OA_L), and high performing older adults (OA_H). Shaded areas around the bars represent the standard error. Each point represents an individual participant.

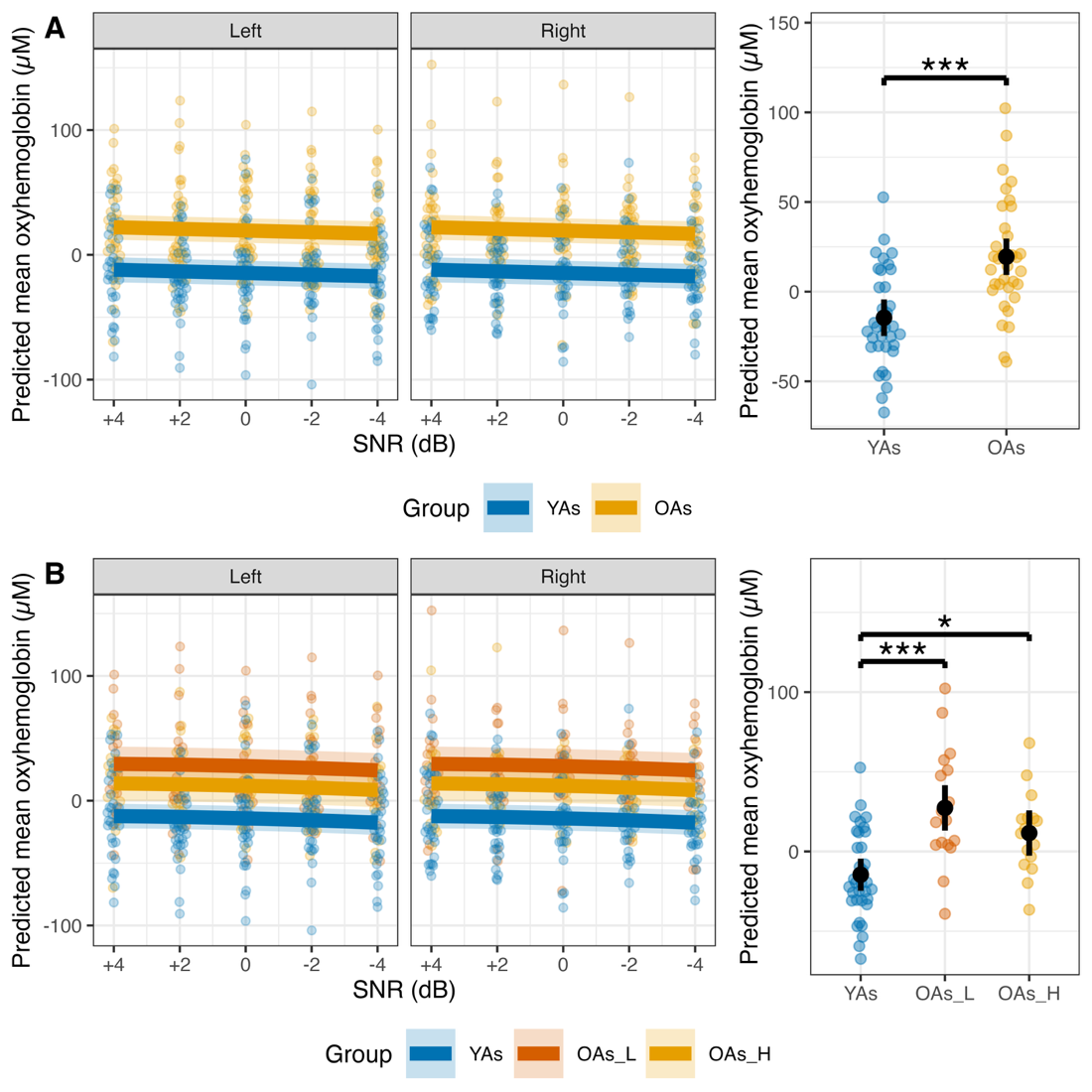

Correlation coefficients by age group and hemisphere. (A) Correlations between accuracy, self-reported listening effort, and HbO within each signal-to-noise ratio (SNR) and hemisphere. (B) Pearson correlations between HbO and demographic measures, including age, best-ear pure tone average (BestPTA), and MoCA scores. Asterisks indicate significant correlations.

*Only a significant correlation was found between DMPFC HbO responses and self-reported listening effort in SNR -4 dB in young adults. All other correlations were not significant.*

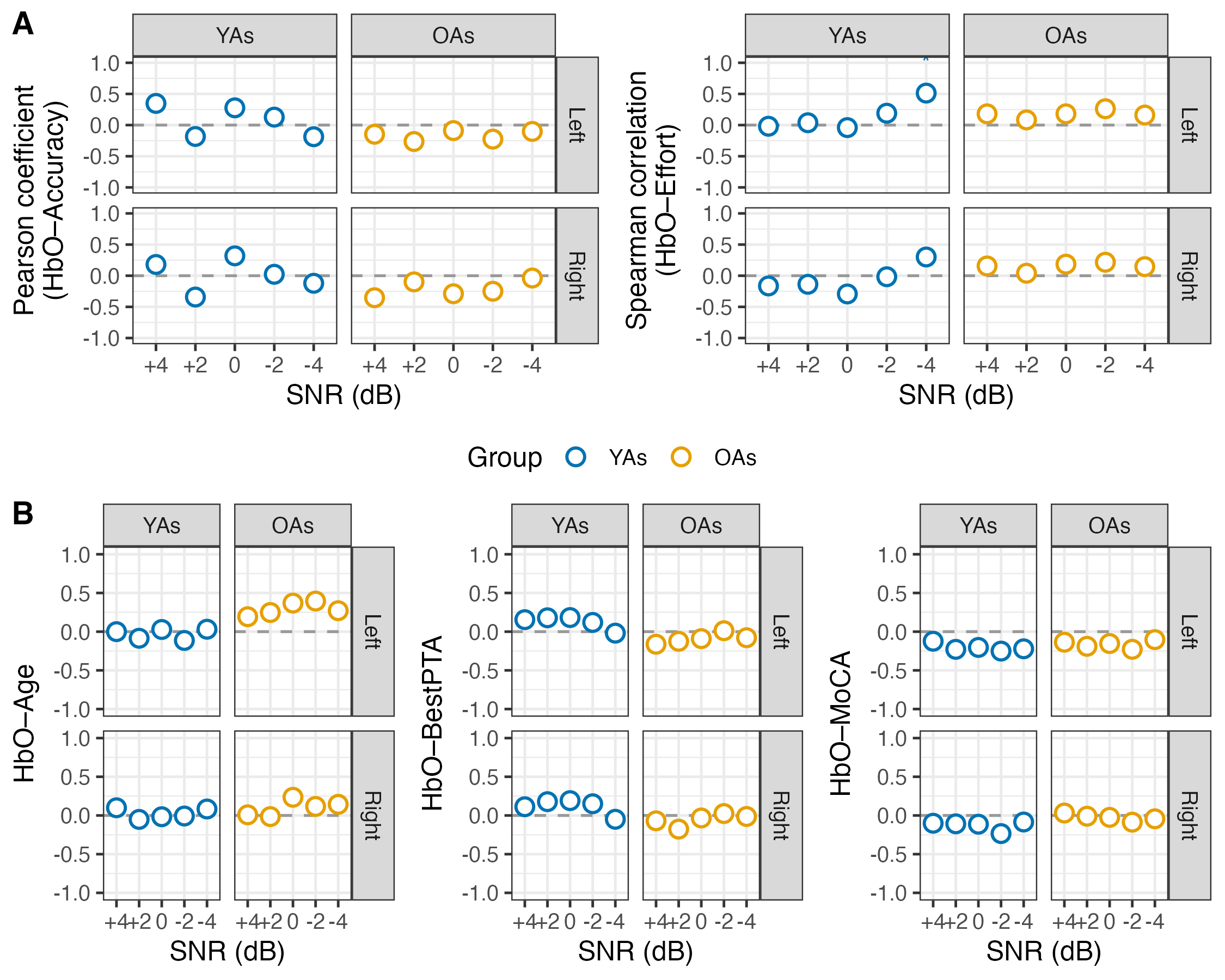
